# Cell type-specific histone acetylation profiling of Alzheimer’s Disease subjects and integration with genetics

**DOI:** 10.1101/2020.03.26.010330

**Authors:** Easwaran Ramamurthy, Gwyneth Welch, Jemmie Cheng, Yixin Yuan, Laura Gunsalus, David A. Bennett, Li-Huei Tsai, Andreas Pfenning

**Affiliations:** Computational Biology Department, Carnegie Mellon University, Pittsburgh PA 15213; Picower Institute for Learning and Memory, Massachusetts Institute of Technology, Cambridge MA 02139; Rush University Medical Center, Chicago IL 60612

## Abstract

We profile genome-wide histone 3 lysine 27 acetylation (H3K27ac) of 3 major brain cell types from hippocampus and dorsolateral prefrontal cortex (dlPFC) of subjects with and without Alzheimer’s Disease (AD). We confirm that single nucleotide polymorphisms (SNPs) associated with late onset AD (LOAD) prefer to reside in the microglial histone acetylome, which varies most strongly with age. We observe acetylation differences associated with AD pathology at 3,598 peaks, predominantly in an oligodendrocyte-enriched population. Strikingly, these differences occur at the promoters of known early onset AD (EOAD) risk genes (*APP, PSEN1, PSEN2, BACE1*), late onset AD (LOAD) risk genes (*BIN1, PICALM, CLU, ADAM10, ADAMTS4, SORL1* and *FERMT2*), and putative enhancers annotated to other genes associated with AD pathology (*MAPT*). More broadly, acetylation differences in the oligodendrocyte-enriched population occur near genes in pathways for central nervous system myelination and oxidative phosphorylation. In most cases, these promoter acetylation differences are associated with differences in transcription in oligodendrocytes. Overall, we reveal deregulation of known and novel pathways in AD and highlight genomic regions as therapeutic targets in oligodendrocytes of hippocampus and dlPFC.

## INTRODUCTION

Alzheimer’s Disease (AD) is the most common age-related neurodegenerative disorder^1^. The hallmarks of AD pathology are numerous and include neuronal loss, synaptic dysfunction, gliosis, and the accumulation of intercellular plaques of amyloid-β (Aβ) protein and intracellular neurofibrillary tangles (NFT) of phosphorylated tau protein (*MAPT*) ^2^.

Aβ plaques are formed by differential proteolytic cleavage of the amyloid β precursor protein (*APP*)^3–6^ by the α-secretase, β-secretase and γ-secretase enzymes^7^. Studies of individuals affected by early onset (<60 yrs.) familial AD (EOAD) have identified causal autosomal dominant mutations primarily in Aβ processing proteins presenilin-1 (*PSEN1*) and presenilin-2 (*PSEN2*), which are part of the γ-secretase complex^8–10^, but also causal mutations or duplications in *APP* itself^11–13^. However, EOAD only accounts for a small minority of AD cases. Late onset sporadic AD (LOAD) is more frequent and accounts for up to 99% or more of AD cases. While increased age is the strongest risk factor and several environmental factors also confer risk for LOAD, its heritability has been estimated to be as high as 79%^14^.

In contrast to EOAD, genetic risk for LOAD is less well understood. The ε4 allele comprising mutations in two codons in Apolipoprotein E (*APOE*) has been identified as the strongest genetic risk factor for LOAD^15–19^. More recently, genome wide association studies (GWAS)^20–27^ have reproduced the *APOE* association and also identified 28 other unique loci harboring genetic variants which increase risk for developing LOAD^26–28^. Strikingly, from the set of most significant (or “sentinel”) single nucleotide polymorphisms (SNPs) derived from GWAS and SNPs in strong linkage disequilibrium (LD) with them, only 2% localize in known exons. Since these SNPs do not alter protein sequence, it is difficult to trace their functional importance in disease onset and progression.

To this end, epigenomic studies are revealing that these SNPs likely alter the function of gene regulatory elements in AD. 26% of these SNPs localize in regions containing promoter histone marks, 69% lie in enhancer states and 46% lie in DNase I accessible sites^26,29^. Further, previous research shows that the human orthologues of enhancers with increased activity in the CK-p25 mouse model of neurodegeneration overlap with non-coding AD associated SNPs^30^. Recently, these SNPs were also found to be primarily contained within microglial enhancers^31^. Furthermore, deregulation of histone 3 lysine 27 acetylation (H3K27ac) and histone 4 lysine 16 acetylation (H4K16ac) was found at loci harboring non-coding AD associated SNPs in the human postmortem AD brain^32,33^. Beyond AD risk loci, changes in histone 3 lysine 9 acetylation (H3K9ac) driven by tau pathology have also been observed in the aging and AD brain^34^.

Gene regulatory elements, especially enhancers, are highly context-specific with differing activities across tissues, cell types and environments^35^. Therefore, it is likely that different cell types in the brain orchestrate different regulatory programs during AD progression. Indeed, many LOAD risk loci are primarily implicated in immune function, suggesting differential AD-associated epigenomic mechanisms in immune cell types such as microglia versus neuronal cell types^30,36–38^. Notably, many of the above-mentioned studies were performed utilizing whole brain tissue, and not all were performed with tissue from AD patients. Therefore, these epigenomic experiments obscure changes that occur within specific cellular populations.

We address these issues by profiling individual cell types deregulated during AD. We utilize fluorescence-activated nuclei sorting (FANS)^39^ to purify neuronal, microglial and other glial populations in the dorsolateral prefrontal cortex (dlPFC) and hippocampus of subjects with and without AD pathology. Then, we perform chromatin immunoprecipitation and sequencing (ChIP-seq) for H3K27ac, which is associated with active promoters and enhancers^40^, to mark putative regulatory elements (peaks) in these populations.

In addition to establishing the first genome-wide H3K27ac profiles in neuronal, microglial, and oligodendrocyte-enriched glial populations from persons with and without AD, our cell type-specific approach confirms enriched H3K27ac signatures at GWAS derived LOAD risk loci primarily in microglia. Further, in both the hippocampus and dlPFC, we find strong Aβ-associated deregulation of H3K27ac in the oligodendrocyte-enriched glial population near AD risk loci and myelin-associated genes. These findings suggest distinct gene-regulatory mechanisms of AD onset and progression in different brain cell types and highlight specific cell types, loci and pathways for future study.

## RESULTS

### Fluorescence-activated nuclei sorting and H3K27ac ChIP-seq of dlPFC and hippocampus

We obtained 10 dlPFC and 16 hippocampus samples from participants in either the Religious Orders Study or Rush Memory and Aging Project (ROSMAP)^41–43^ (mean age = 87.84, s.d.=7.75, range=74.77-101.94). 5 of 10 dlPFC samples and 10 of 16 hippocampus samples displayed high Aβ load across the brain, indicative of LOAD (mean percentage area occupied by Aβ across 8 brain regions = 7.30, s.d = 4.14, range = 2.31-15.40) (**Supplementary Table 1, Supplementary Figure 1**). The brains with Aβ load also displayed high overall neurofibrillary tangle density (mean density of NFT across 8 brain regions = 22.81, s.d. = 13.73, range = 1.80-61.01). The self-reported sex of 6 of the 10 dlPFC samples was male, and the remaining 4 were female. Of the 16 hippocampus samples, the self-reported sex of 6 was male, and the remaining 10 were female.

For each sample, we used FANS to collect NeuN+, Pu.1+, and NeuN-/Pu.1-nuclei to obtain putative neuronal, microglial, and other glial populations, respectively (**Figure 1a, Supplementary Figure 2**)^39^. On each collected population, we performed ChIP-Seq for H3K27ac, which is associated with transcriptionally active promoters and enhancers^40^. We assessed sample quality by calling regions of H3K27ac enrichment (peaks) for each individual sequencing sample and computing quality metrics based on standard ENCODE guidelines^44^. We detected an average of 91,614 (s.d = 21,197, range=50,662-149,681) peaks per sample. These peaks overlapped with a large fraction of the sequencing reads (mean FRiP = 0.256, s.d. = 0.136, range=0.047-0.567), comparable to previous high quality H3K27ac profiles^35^. We curated samples further based on normalized strand cross correlation (NSC) and relative strand cross correlation (RSC) measures to ensure that we retained the highest quality sequencing samples for all downstream analysis (**Methods, Supplementary Figure 1**).

**Figure 1:**
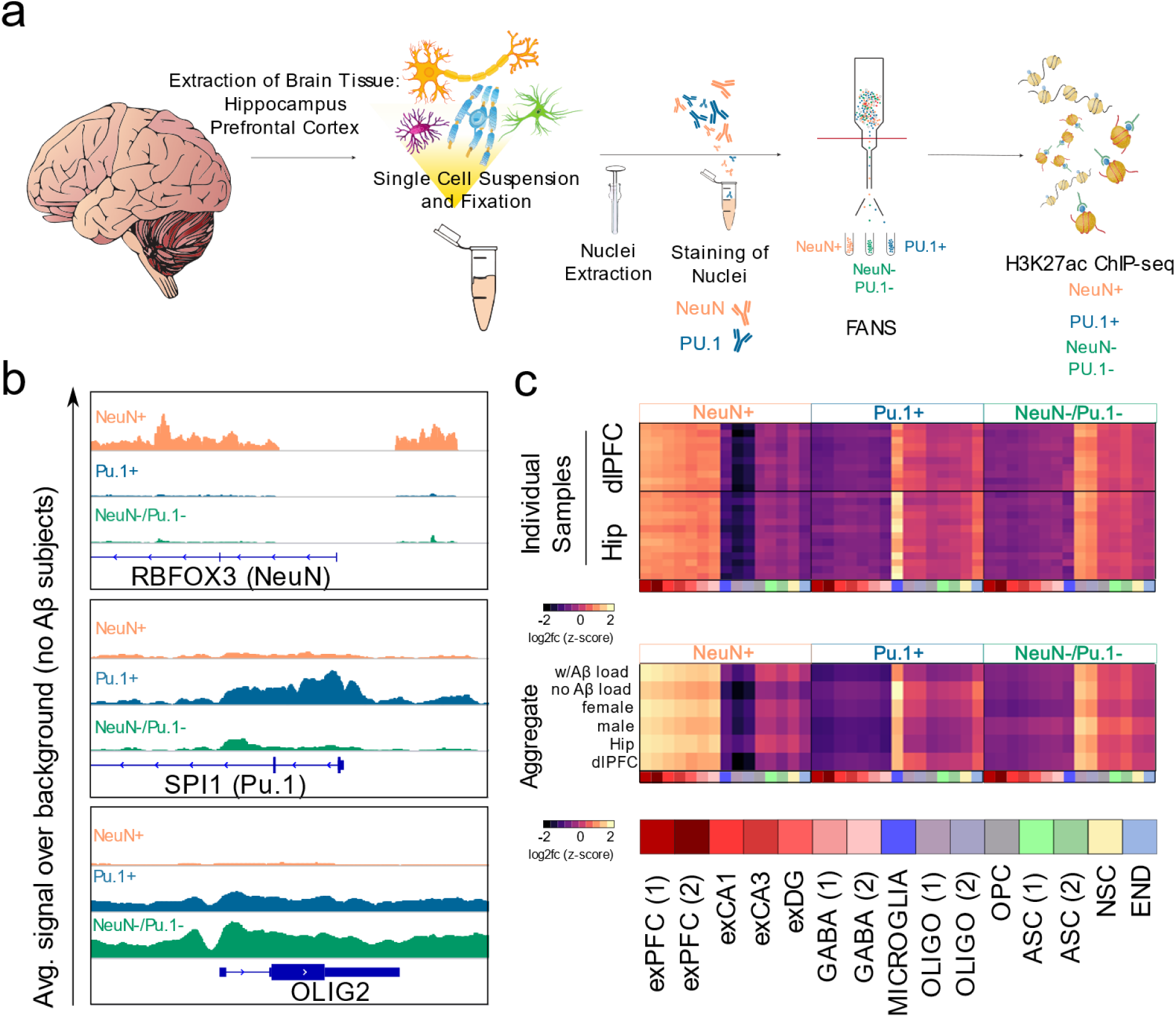
FANS sorting captures neurons, microglia and oligodendrocyte enriched populations from postmortem brain tissue. **a**. Workflow for sorting nuclei and performing H3K27ac ChIP-seq from postmortem human brain tissue **b**. H3K27ac signal over background (Input) averaged across subjects without Aβ load for each of the three populations near RBFOX3 (NeuN), SPI1 (Pu.1) and OLIG2 (an oligodendrocyte marker) **c. Top**. for every individual hippocampus and dlPFC tissue sample, fold-change (log2 transformed) of H3K27ac signal in each focal cell population over the other two non-focal cell populations, averaged across peaks near the promoters (<5kb from transcription start site) of genes defined to be markers for 15 different cell types in Habib et al^46^. **bottom**. collapsed versions of top heatmap representing averages across subjects defined by different stratifications of Aβ load, sex and brain region. Abbreviated labels for the single nucleus analysis clusters are presented. exPFC, glutamatergic neurons from the PFC; GABA, GABAergic interneurons; exCA1/3, pyramidal neurons from the hippocampus CA region; exDG, granule neurons from the hippocampus dentate gyrus region; ASC, astrocytes; MICROGLIA, microglia; OLIGO, oligodendrocytes; OPC, oligodendrocyte precursor cells; NSC, neuronal stem cells; END, endothelial cells.

Then, for each brain region and each cell population, we used ENCODE recommended approaches^44^ to call H3K27ac peaks that are reproducible across subjects with Aβ load, and separately, peaks that are reproducible across subjects without Aβ load. We created a union of all these peak sets representing the combined histone acetylome of the three profiled cell populations in the dlPFC and hippocampus of subjects with and without Aβ load (**Supplementary Table 2**). We then used DESeq2^45^ to obtain peaks that are significantly hyperacetylated in (i) the NeuN+ population relative to the Pu.1+ and NeuN-/Pu.1-populations, (ii) the Pu.1+ population relative to the NeuN+ and NeuN-/Pu.1-populations, and (iii) the NeuN-/Pu.1-population relative to NeuN+ and Pu.1+ populations (FDR q<0.05). We performed principal component analysis (PCA) of all samples and observed groupings primarily based on FANS population, with 53% of the variance explaining the difference between NeuN+ samples and other samples (**Supplementary Figure 3**).

### Active promoters and enhancers in neurons, microglia and oligodendrocyte enriched glia

As a first step to assess the efficacy of FANS sorting, we generated genome browser tracks of H3K27ac signal for each population by averaging signal across control subjects displaying no Aβ. We visualized these genome browser tracks near genes encoding the cell type-specific proteins used to sort out neurons and microglia – *RBFOX3* which encodes NeuN, and *SPI1* which encodes Pu.1 (**Figure 1b**). As expected, we observed average hyperacetylation at the locus containing RBFOX3 in the NeuN+ samples and average hyperacetylation in the Pu.1+ samples at the locus containing *SPI1*, suggesting successful sorting. Interestingly, we observed hyperacetylation in the NeuN-/Pu.1-samples near genes that are highly expressed in oligodendrocytes, such as *OLIG2*, suggesting oligodendrocyte enrichment.

To confirm these initial qualitative assessments of sorting efficacy, and to identify the cell types captured in the NeuN-/Pu.1- population, we performed a more rigorous comparison of our H3K27ac ChIP-seq data with an independent higher-resolution single nucleus gene expression (snRNA-seq) dataset from human prefrontal cortex and hippocampus described in Habib et al.^46^. As expected, the NeuN+ samples displayed significant hyperacetylation on average at peaks annotated to nearby genes significantly upregulated in excitatory neuron clusters from prefrontal cortex (adjusted p=1.8e-204, 1.25e-92), hippocampus (adjusted p=3.37e-173, 1.20e-80), and dentate gyrus (adjusted p=2.23e-35), and also GABAergic neuron clusters (adjusted p=5.4e-28, 1.4e-31) (**Supplementary Figure 4b**). Similarly, the Pu.1+ samples displayed significant hyperacetylation on average at peaks annotated to genes significantly upregulated in microglia (adjusted p = 3.18e-22). Strikingly, the NeuN-/Pu.1-samples displayed significant hyperacetylation on average at peaks annotated to genes significantly upregulated in oligodendrocyte clusters (adjusted p= 1e-58, 1.46e-25), but not any of the other cell types queried, confirming oligodendrocyte enrichment.

Since AD pathology, brain region, and sex could potentially influence sample quality and sorting efficacy, we repeated this analysis separately for (i) samples with and without Aβ, (ii) samples from dlPFC and hippocampus, (iii) male and female samples, and (iv) each sample individually. In each of these analyses, we observed neuronal enrichment in NeuN+ samples, microglial enrichment in Pu.1+ samples, and oligodendrocyte enrichment in NeuN-/Pu.1-samples (**Figure 1c, Supplementary Figure 4c**). Since enhancers are known to have long range effects and may not necessarily regulate their nearest genes, we also restricted the analysis to peaks proximal to gene transcription start sites (TSS) (<5 kilobases) and observed the same results (**Figure 1c, Supplementary Figure 4a**). Therefore, we conclude that the NeuN+ population successfully captures neurons, the Pu.1+ population successfully captures microglia, and the NeuN-/Pu.1-population is highly enriched for oligodendrocytes.

Together, our peak annotations represent the first genome-wide maps of H3K27ac in microglia, neurons, and oligodendrocyte-enriched glial (OEG) populations in the human hippocampus and dlPFC of subjects with and without Aβ. These annotations enable a better understanding of the gene regulatory roles of the profiled cell types in many different contexts, not limited to AD. Nevertheless, in the next sections, we utilize these annotations to understand cell type-specific epigenomic mechanisms in AD. First, we compare these annotations with GWAS data to annotate LOAD associated SNPs to the cell types and regulatory elements they may potentially disrupt. Second, we perform multiple histone acetylome-wide association studies in each sex, brain region, and cell type to identify AD-associated variations in acetylation. Third, we perform a histone acetylome-wide association study to identify acetylation differences associated with age in each cell type.

### GWAS derived common SNPs associated with LOAD risk preferentially colocalize with the microglial histone acetylome

We performed partitioned heritability analysis by stratified LD score regression^47–49^ (S-LDSC) to estimate the strength of colocalization between H3K27ac peaks that are significantly hyperacetylated on average across subjects in the 3 populations and AD SNP heritability derived from two large AD GWAS meta analyses (Jansen et al. and Kunkle et al.)^26,27^. Strikingly, microglial hyperacetylated peaks displayed a statistically significant preference for colocalization with AD SNP heritability (**Figure 2a and b;** Jansen et al. GWAS coefficient = 1.6e-08, p = 5.28e-5, Kunkle et al. GWAS coefficient = 1.94e-08, p = 3.74e-3) relative to neuronal and OEG hyperacetylated peaks. Since choice of computational method can influence these assessments, we repeated the analysis with an independent method that utilizes a permutation test^35,50^. We again observed that AD SNP heritability has a strong preference for colocalization with microglial hyperacetylated peaks (**Supplementary Figure 5;** Kunkle log2FC= +0.39, adjusted p=1e-06, Jansen log2FC=+0.33, adjusted p=1e-06). Further, conducting these analyses with reproducible peaks for each cell type, as opposed to hyperacetylated peaks led to similar results. These findings agree with previous analyses conducted on myeloid cells^50–52^, reinforcing the hypothesis that myeloid cell gene regulation strongly influences predisposition towards AD.

**Figure 2:**
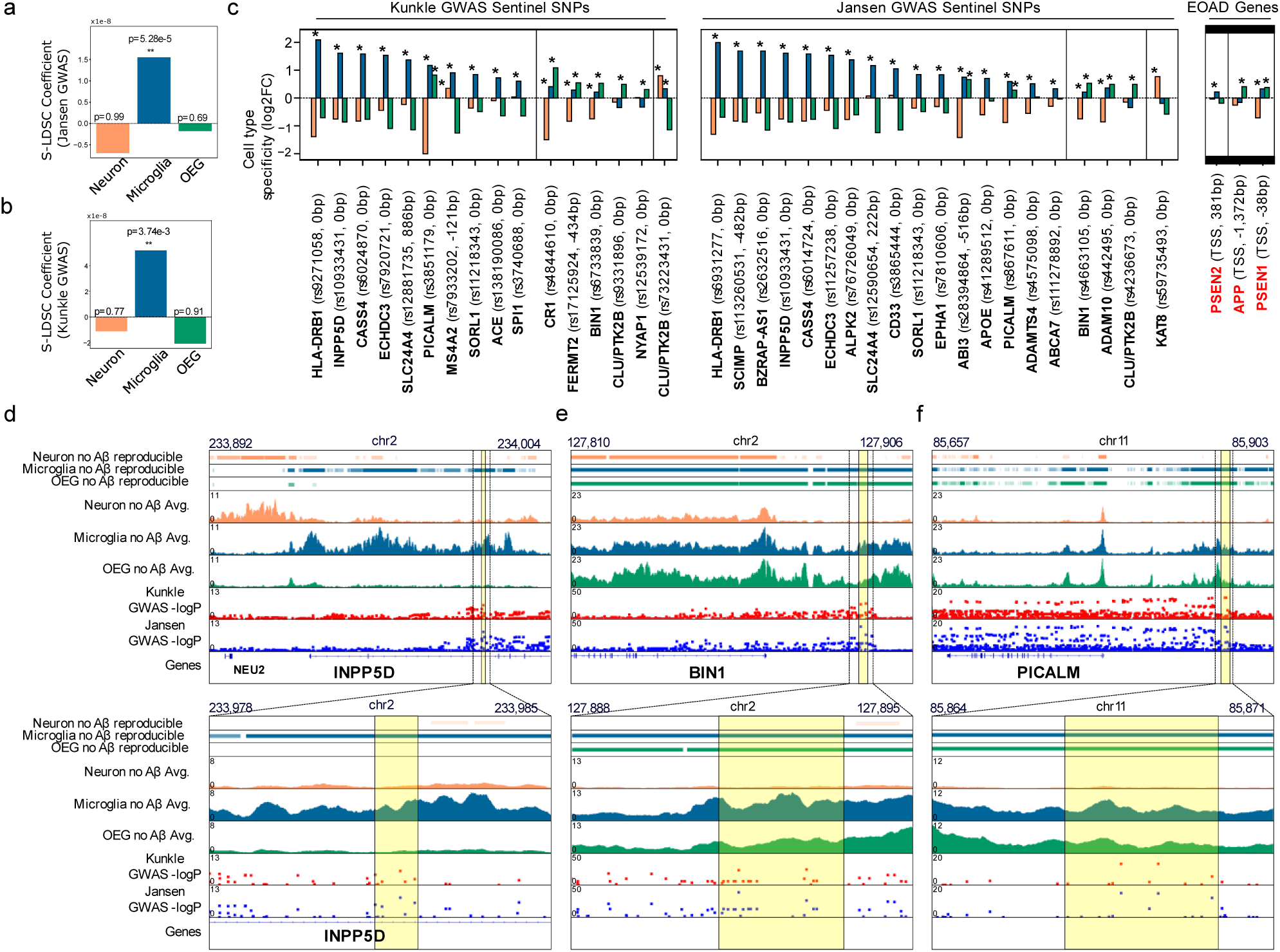
AD associated SNPs derived from GWAS prefer to colocalize with peaks enriched in the microglial population relative to peaks enriched in the OEG and neuronal population. **a and b**. Results of stratified LD score regression on two large AD GWAS studies (Jansen et al and Kunkle et al)^26,27^ on hyperacetylated peaks in each population. Plot shows the estimated LD score regression coefficient for the three peak sets. p-values are indicated above each bar. **c**. Cell type-specificity of peaks annotated to known sentinel non-synonymous SNPs at AD risk loci identified by Jansen and Kunkle et al. Plot shows fold change (log2-transformed) in H3K27ac signal for each population against the other two populations for (i) in black: peaks closest to the sentinel SNP at each locus associated with AD from GWAS, and (ii) in red: promoter peaks of early onset AD risk genes (*APP, PSEN1, PSEN2*). *indicates DeSeq2 FDR q<0.05. Sentinel SNPs where the closest SNP is >1kb away are not included **d**,**e and f**. Genome browser tracks of (i) reproducible peaks in each cell type, (ii) average H3K27ac signal in subjects without Aβ load for each cell type, and (iii) Manhattan plots of Jansen et al and Kunkle et al. GWAS studies at loci where sentinel non-coding SNPs overlap peaks enriched in non-neuronal cell types; at the *INPP5D* containing locus, the sentinel SNP rs10933431 overlaps a peak that is active only in microglial population but not OEG and neuronal populations; at the locus containing *BIN1*, the top two AD-associated SNPs based on GWAS p-value, rs4663105 and rs6733839 overlap peaks active in the microglial and OEG populations but not in the neuronal population; at the locus containing *PICALM*, the top two SNPs, rs10792832 and rs3851179 also overlap non-neuronal enhancers. Region of overlap highlighted by yellow box.

We note that neuronal hyperacetylated peaks overlap with a lower number of GWAS derived AD associated SNPs compared to microglial and OEG hyperacetylated peaks (**Supplementary Figure 5; Figure 2c**). This finding is consistent with previous analyses conducted on bulk brain tissue maps of histone modifications^35,50^ and open chromatin^37,38^, where signal from neuronal regulatory elements is dominant. Since biases in GWAS sampling and neuronal sample quality could potentially influence the results of these analyses, we used S-LDSC to partition Schizophrenia SNP heritability^53^ across the hyperacetylated peaks in the 3 populations. Only neuronal hyperacetylated peaks displayed significant colocalization (**Supplementary Figure 6;** coefficient = 1.5e-07, p=1.4e-8). This agrees with previous findings about Schizophrenia^54,55^, and therefore, confirms that the analysis is robust to biases in GWAS sampling and neuronal sample quality.

### Interpreting cell-type specificity and potential disruptions of non-coding AD associated variants

We annotate non-coding sentinel SNPs identified in Jansen et al. and Kunkle et al. to nearby peaks (<1kb cutoff), enabling assessment of their potential cell type-specific functional effects (**Figure 2c, Table 1**). As expected, at a majority of GWAS derived risk loci, the sentinel SNPs directly overlap with peaks that are most strongly hyperacetylated in microglia. However, many sentinel SNPs including SNPs at loci containing *BIN1, CLU, ADAM10*, and *CR1* directly overlap peaks that are most strongly hyperacetylated in OEG. Only 2 sentinel SNPs overlap with peaks that are most strongly acetylated in neurons. Further, the peaks closest to the TSS of *APP* and *PSEN1* display the strongest acetylation in OEG, whereas the peak closest to the TSS of *PSEN2* display the strongest acetylation in microglia.

Overall, these annotations improve the interpretation of the functional effects of non-coding LOAD-associated SNPs. We point out specific examples such as the locus containing the *INPP5D* gene, where the sentinel SNP rs10933431 (GWAS p-values = 8.9e-10, 2.5e-07) overlaps a peak that is acetylated only in microglia but not neurons and OEG ((**Figure 2d**). Previously, rs10933431 has been shown to overlap with DNase I hypersensitive sites in peripheral blood cells and tissues, including natural killer cells and CD14+ monocytes^29^. Further, rs10933431 disrupts a binding motif for the paired box transcription factor Pax-5^29,56^, which is important for immune cell maturation. Combined, this suggests that rs10933431 is likely altering regulatory function in immune cell types and microglia, and future studies on the functional effect of this SNP should include culture or model systems that can capture phenotypes of these cell types.

Secondly, at the locus containing the *BIN1* gene, which displays the second largest genome wide AD association behind the *APOE* containing locus, two sentinel SNPs overlap a peak which is acetylated in both microglia and OEG (**Figure 2e**), but not neurons. One of the SNPs, rs4663105 (GWAS p-values = 3.37e-44, 2.16e-26) has known expression quantitative loci (eQTL) associations with *BIN1* gene expression in whole blood and lymphoblastoid cells^57,58^. Similarly, the other SNP, rs6733839 (GWAS p-values = 1.28e-29, 4.02e-28) is a *BIN1* eQTL in artery and lymphoblastoid cells^57,58^. Previously, rs6733839 has been shown to overlap with DNase I hypersensitive sites in natural killer cells and CD14+ monocytes^29^. Recently, another study has found that the enhancer overlapping rs6733839 interacts with the *BIN1* promoter in microglia^31^. Further, deletion of this enhancer using CRISPR-Cas9 editing altered *BIN1* expression in inducible pluripotent stem cell (iPSC) derived microglia, but not neurons and astrocytes. This points towards a role for rs6733839 in disrupting *BIN1* expression in cells of the myeloid lineage. However, effects of rs6733839 on *BIN1* expression in oligodendrocytes have not been previously studied and therefore, cannot be excluded since the peak is also strongly acetylated in OEG. Further, the other sentinel SNP, rs4663105, could also potentially exert effects on *BIN1* expression in microglia or oligodendrocytes, and future studies can help clarify this.

Similarly, at the locus containing *PICALM*, one sentinel SNP, rs10792832 (GWAS p-values = 7.36e-18, 7.55e-16) and another SNP in tight linkage, rs3851179 (GWAS p-values = 2.02e-17, 5.81e-16) overlap non-neuronal peaks (**Figure 2f**). This suggests that these SNPs are potentially exerting effects on expression of *PICALM* in microglia and/or oligodendrocytes, and models of these cell types should be included in future studies to assess their functional effects.

These examples highlight the utility of our data resource in informing future studies of non-coding SNPs associated with traits that include, but are not limited to AD.

### OEG display strongest AD associated acetylation differences

In each brain region, sex, and cell type, we used a histone acetylome-wide association study to identify acetylation differences between subjects with and without Aβ load using DESeq2^45^. Overall, we discovered 3598 amyloid-associated differentially acetylated regions (DARs) across all experiments (**Supplementary Table 3, Figure 3a**).

**Figure 3:**
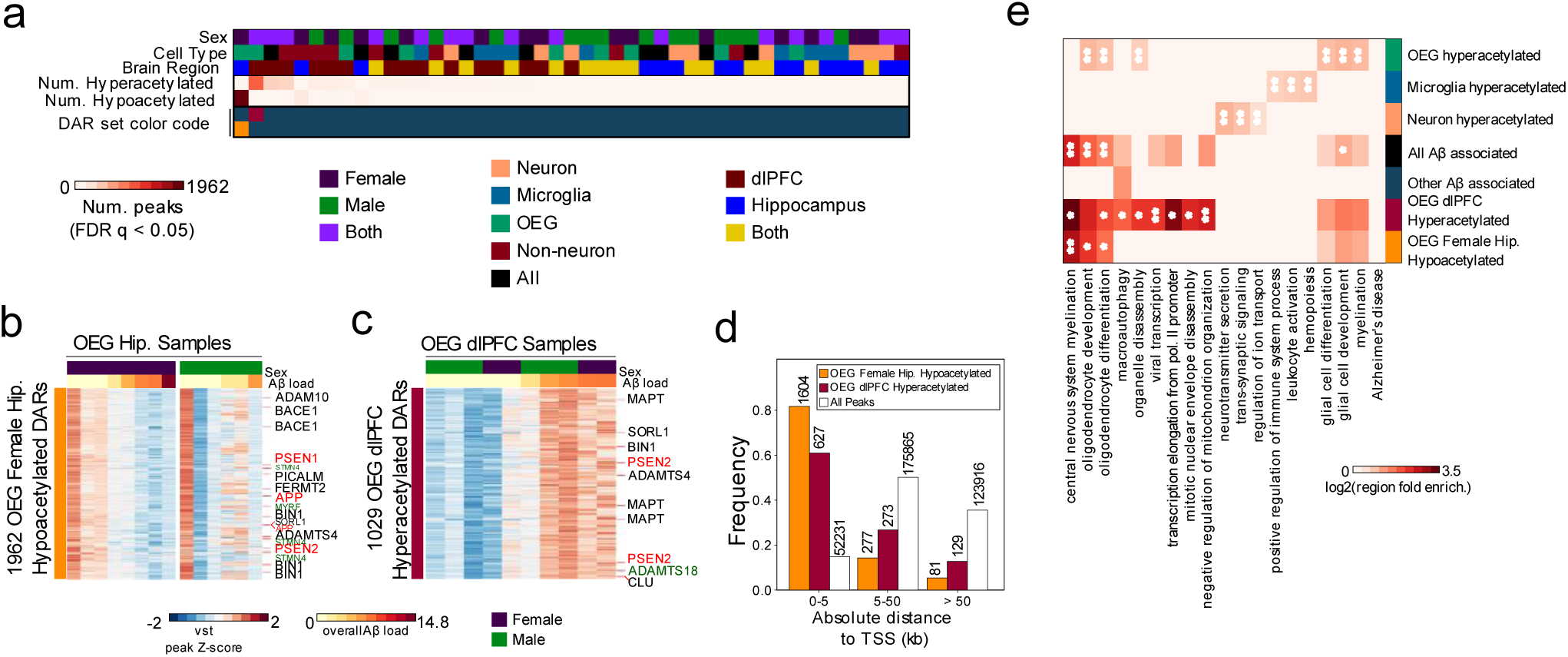
OEG display the strongest acetylation differences associated with Aβ load which includes peaks annotated to known genes associated with EOAD and LOAD risk. **a**. heatmap displaying number of significantly hyperacetylated (log2fc>0, FDR q<0.05) and significantly hypoacetyled peaks (log2fc<0, FDR q<0.05) in each stratification of brain region, sex and cell type that was tested **b**. Heatmap showing normalized acetylation levels at 1962 AD hypoacetylated DARs discovered in female hippocampus OEG samples, rows represent the 1962 DARs and columns represent OEG samples from the hippocampus. Measured Aβ load for each sample is indicated at the bottom of the heatmap, a heatmap for male hippocampal glia samples is also included for comparison, peaks annotated to known EOAD and LOAD risk are labeled in red, peaks near *STMN4* (highest hypoacetylation) and *MYRF* are annotated in black **c**. heatmap of 1029 DARs displaying hyperacetylation in OEG in dlPFC samples. Peaks annotated to known EOAD and LOAD risk genes are labeled in red, the promoter peak at *ADAMTS18* (highest hyperacetylation) is annotated in black **d**. distance to TSS distribution of i) 1962 OEG female hippocampus hypoacetylated, ii) 1029 OEG dlPFC hyperacetylated DARs and iii) the full set of peaks **e**. heatmap showing enrichment of top gene ontology terms for 6 peak sets 1) 1962 OEG female hippocampus hypoacetylated 2) 1029 OEG dlPFC hyperacetylated 3) all other Aβ associated DARs 4),5), and 6) neuron, microglia and OEG cell type-specific hyperacetylated peaks. Color intensity represents hypergeometric fold enrichment in number of peaks, * indicates FDR q<0.05, ** indicates FDR q<0.01

Unexpectedly, we observed minimal differences in acetylation associated with Aβ load in microglia. In contrast, the OEG population is associated with the largest acetylation differences and contributes to a majority of identified DARs. We discovered two DAR sets, the largest in female hippocampus OEG samples (1962 hypoacetylated; q<0.05) and the second largest in dlPFC OEG samples (1029 hyperacetylated; q<0.05) that make up 80.3% (2,890) of the full set of 3,598 DARs. We confirmed that both DAR sets display progressive trends of differential acetylation when treating Aβ load as a continuous variable. Further, in a post-hoc analysis, we controlled for covariates such as age at death and years of education which display no correlation with acetylation levels at these DARs in the corresponding OEG populations (**Supplementary Figure 7**).

### Hypoacetylation in OEG of the hippocampus

We discovered 1962 hypoacetylated DARs in female hippocampus OEG samples, 81.7% of which are peaks proximal to TSS (<5kb) (hypergeometric test p-value=0, **Figure 3d**), suggesting strong links with promoter activity and gene transcription. Strikingly, this hypoacetylated DAR set includes peaks at the promoters of *APP, PSEN1*, and *PSEN2*, the three genes associated with EOAD risk, as well as promoters at several LOAD risk loci identified by GWAS, including *BIN1, PICALM, ADAMTS4, ADAM10*, and *FERMT2* (**Supplementary Table 4, Figure 3b, Figure 4a-i, Supplementary Figure 8**). Notably, promoters of genes involved in all three secretase complexes including α-secretase (*ADAM10*), β-secretase (*BACE1*), and γ-secretase (*PSEN1, PSEN2* and *PSENEN*) are hypoacetylated, suggesting Aβ processing is directly disrupted in oligodendrocytes.

**Figure 4:**
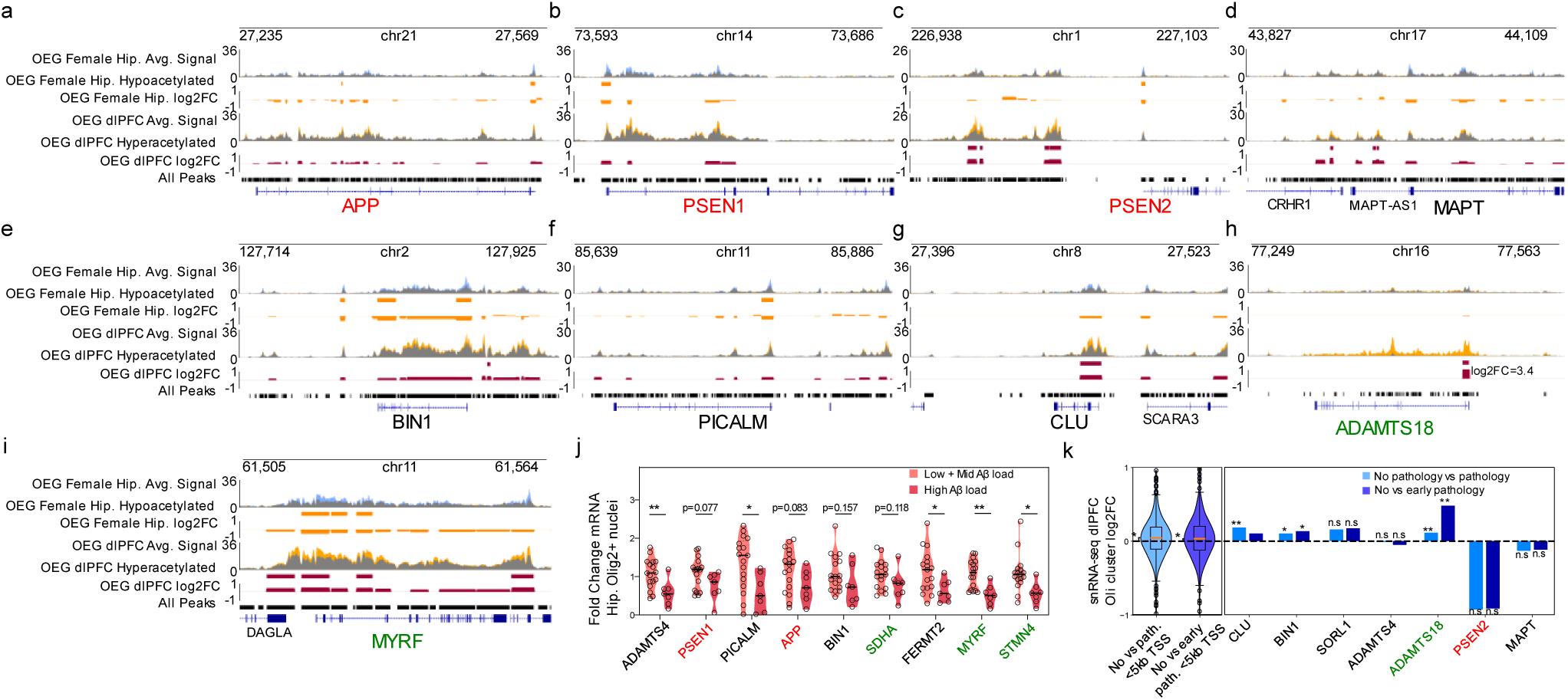
EOAD and LOAD may share common epigenomic perturbations and pathogenic mechanisms in oligodendrocytes. **a-i**. genome browser tracks displaying average signal in OEG samples of subjects with (yellow) and without Aβ load (blue) corresponding to two biggest DAR sets at known EOAD and LOAD risk loci, as well as strongly differentially acetylated regions near *ADAMTS18* and *MYRF* **j**. qRT-PCR in hippocampal Olig2+ nuclei from a larger cohort of subjects reveals an associated decrease in transcription near hypoacetylated regions included in the female hippocampal OEG DAR set **k**. comparison with existing snRNA-seq from dlPFC of subjects from the same cohort reveals an average increase in gene expression near hyperacetylated regions in OEG dlPFC. Further, specific genes associated with AD risk and displaying high hyperacetylation nearby (*ADAMTS18*) display an increase in transcription in AD.

We performed gene ontology analysis of these DARs using GREAT^59^ which revealed an enrichment for central nervous system myelination, oligodendrocyte development, and oligodendrocyte differentiation (**Supplementary Table 5, Figure 3e**). We also observed hypoacetylation near genes encoding the five mitochondrial complexes that regulate oxidative phosphorylation (**Supplementary Figure 9**). Since acetylation differences associated with myelination, oligodendrocyte differentiation and oxidative phosphorylation occurs in tandem with acetylation differences at AD risk genes and amyloid processing, these pathways may directly contribute to AD onset and progression.

To assess whether these acetylation differences are associated with differences in transcription in oligodendrocytes, we performed quantitative RT-PCR (qRT-PCR) for multiple genes annotated to peaks in this DAR set in oligodendrocyte (Olig2+) nuclei collected from hippocampus samples of a larger set of subjects with and without Aβ from the same cohort (**Figure 4j, Supplementary Figure 10**). *ADAMTS4, PICALM*, and *FERMT2* displayed significant decreases in transcript levels when comparing low and mid-Aβ load subjects against high Aβ load subjects. *APP* (p=0.083), *BIN1* (p=0.157), and *PSEN1* (p=0.077) displayed similar fold decreases that did not meet the p-value cutoff. Transcriptional differences did not display sex-specificity. Combined, this strongly suggests that EOAD and LOAD may share common pathogenic mechanisms in the oligodendrocytes of the human hippocampus.

We discovered the strongest hypoacetylation in this DAR set at a peak annotated to the *STMN4* gene (log2FC=-1.12, FDR q=1e-6) which is preferentially expressed in brain tissue^60^ and has known functions in neuron projection development and microtubule polymerization^61^. Notably, several other peaks near the *STMN4* gene, including a peak at the *STMN4* promoter, displayed significant hypoacetylation. *MYRF*, a transcription factor which directly activates myelination^62^ and has been previously linked to LOAD risk^63^, also displayed strong promoter hypoacetylation (log2FC=-0.48, FDR q=0.03). *STMN4* and *MYRF* also display significantly reduced transcription in qRT-PCR analysis of oligodendrocyte in subjects with AD. We highlight these myelination-associated genes as high-confidence targets for further investigation in neurodegenerative disorders.

While peaks in this DAR set are annotated to loci associated with AD risk, we did not observe significant colocalization of this DAR set with GWAS derived AD-associated SNPs relative to the full set of peaks active in the profiled cell types and brain regions of AD and non-AD subjects (Jansen coefficient =3.9e-08, p=0.198, Kunkle coefficient=1.11e-07, p=0.29). Therefore, SNPs associated with AD from GWAS are unlikely to directly alter the regulatory function of these DARs directly.

### Hyperacetylation in OEG of the dlPFC

We discovered the second largest histone acetylome variation comprising 1029 hyperacetylated DARs in dlPFC oligodendrocyte-enriched glia (OEG) samples. While this DAR set is distinct from the DARs discovered in female hippocampus OEG samples, and contains a lower proportion (60.9%) of TSS proximal peaks (<5kb), we again observed significant hyperacetylation at both EOAD and LOAD risk loci (**Supplementary Table 4, Figure 3c, Figure 4a-i, Supplementary Figure 8**). This includes four distal intergenic peaks annotated to *PSEN2*, one distal peak annotated to *BIN1*, and peaks overlapping the promoters of *CLU, ADAMTS4*, and *SORL1*. Furthermore, we observed significant hyperacetylation at three distal peaks annotated to the *MAPT* gene, which encodes for the tau protein, involved in formation of NFTs.

We performed gene ontology enrichment analysis of this DAR set which again revealed a strong enrichment for central nervous system myelination and oligodendrocyte differentiation (**Figure 3e, Supplementary Table 5)**. In addition, we discovered enrichment for mitochondrion organization, macroautophagy and viral transcription.

We tested whether these acetylation differences are associated with differences in transcription in dlPFC oligodendrocytes by comparing with a previously published snRNA-seq study of dlPFC in AD^64^. On average, genes annotated to these DARs display higher transcription levels in oligodendrocytes of subjects with AD (**Figure 4k**) compared to subjects without AD. Genes associated with LOAD risk including *CLU* and *BIN1* displayed statistically significant upregulation, while *SORL1* (FDR q no vs path.=0.26, FDR q no vs early path.=0.70) displayed upregulation that did not meet the q-value cutoff. We note that *PSEN2* (FDR q no vs path.=0.18, FDR q no vs early path.=0.21) and *MAPT* (FDR q no vs path.=0.1, FDR q no vs early path.=0.09) display a downregulation in transcription with AD pathology, which did not meet the q-value cutoff. Hyperacetylated peaks near *PSEN2* and *MAPT* are distal to the TSS, and therefore, are probably enhancer peaks. Enhancers have been known to regulate target gene expression over long distances and hence, effects on distal transcripts cannot be excluded.

We observed the strongest hyperacetylation at a peak near the promoter of the *ADAMTS18* gene (log2FC=3.4, FDR q=5.1e-81), which is a member of the *ADAMTS* family of metalloproteinases with thrombospondin motifs. This family of proteins is known to play a role in neuroplasticity and has been widely studied for its role in AD^65^. Overall, 38 different peaks annotated to the *ADAMTS18* gene displayed significant hyperacetylation across DAR sets specific to dlPFC microglia and oligodendrocyte-enriched glia, but not neurons. *ADAMTS18* also displayed statistically significant increase in transcription in dlPFC oligodendrocytes of AD subjects. These results suggest that *ADAMTS18* gene regulation is heavily altered in AD in the dlPFC and resides in an important oligodendrocyte and microglia-specific locus that requires further study.

Similar to the female hippocampus OEG hypoacetylated DAR set, we did not observe significant colocalization between GWAS derived AD-associated SNPs and peaks in this DAR set using S-LDSC (Jansen GWAS coefficient =-2.57e-08, p=0.70, Kunkle GWAS coefficient=-1.26e-07, p=0.74). Therefore, SNPs associated with AD from GWAS are unlikely to alter the regulatory function of these DARs directly.

Overall, we reveal that common pathways associated with both early and late onset AD are likely perturbed at the epigenomic level in oligodendrocyte-enriched glia. We show that amyloid processing, central nervous system myelination and oligodendrocyte processes are significantly altered in dlPFC and hippocampus of subjects with amyloid pathology and display acetylation differences in tandem with AD risk genes. We also highlight novel genomic loci that display large changes in acetylation in glia in AD including *ADAMTS18, STMN4* and *MYRF*. Taken together, the sets of DARs we have described are strong candidate targets for AD therapeutics in oligodendrocytes that utilize technologies such as CRISPR-Cas9 genome editing^66^.

We also performed unsupervised clustering of the full set of 3,598 DARs to identify modules that display correlated acetylation across the profiled cell type populations and brain regions. We separately clustered TSS proximal and TSS distal DARs to identify putative promoter and putative enhancer modules, respectively. Modules of both proximal and distal peaks tended to separate based on cell type-specificity. We note that majority of peaks display microglia or OEG specificity (**Supplementary Figure 11**) with little or no acetylation in neuronal samples. This highlights the utility of our data resource in identifying novel gene regulatory modules that are cell type-specific and associated with disease.

### Age associated acetylation differences are enriched in the microglial population

While microglial H3K27ac displays strong colocalization with GWAS derived AD associated SNPs, unexpectedly, the microglial population displays very few acetylation differences associated with Aβ load. Contrastingly, the microglial population displays the strongest age-associated acetylation differences encompassing both dlPFC and hippocampus, in an analysis that controlled for Aβ load, sex, and brain region differences (**Supplementary Figure 12**). We identified 391 peaks that are significantly hypoacetylated with increasing age and 53 peaks that are significantly hyperacetylated with increasing age (FDR q < 0.05) (**Supplementary Table 6**). We mapped these peaks using GREAT^59^ and discovered 2 hypoacetylated peaks annotated to the amyloid precursor protein (*APP*) gene, and 6 hypoacetylated peaks near the *LRRTM3* gene, which is involved in positive regulation of Aβ formation (**Supplementary Table 7**). This suggests that Aβ processing may be altered in microglia with increasing age. We also observed hyperacetylation at 3 distal peaks annotated to the FKBP4 gene, which is involved in tau protein binding and influences neurofibrillary tangle formation. Although further investigation is required, these findings point towards a role for age-associated epigenomic changes in microglia influencing the onset and progression of LOAD.

## DISCUSSION

We report the first H3K27ac maps for sorted neurons, microglia, and oligodendrocyte-enriched glia from both the hippocampus and dlPFC of postmortem human brain tissue. We find microglial H3K27ac peaks have a stronger preference for colocalization with common SNPs associated with LOAD risk relative to the other neural cell types profiled, supporting previous findings^30,31,37,38^. While this suggests a significant causal role for LOAD risk loci influencing AD predisposition and progression through microglial processes, perhaps unexpectedly, comparison of H3K27ac peaks by AD diagnosis in microglia revealed few differences. Instead, we report H3K27ac is altered significantly with age in microglia, leading us to conclude that amongst the individuals analyzed, microglial H3K27ac is more responsive to advances in age than to Aβ load. We note that heterogeneity within the microglial population in disease has been previously reported^52,67^ and therefore, the possibility of AD associated gene regulatory differences in microglia cannot be excluded based on our study which profiles the microglial population in bulk, and hence, represents average microglial signal. However, recent single cell transcriptome profiling of microglia in AD subjects revealed no differences in both the composition of microglial subpopulations as well as gene expression, supporting the findings from our study^68^.

Beyond microglia, we also find a subset of AD risk loci have significant H3K27ac signal in oligodendrocyte-enriched glia relative to other cell types. These include risk loci associated with genes *CLU, BIN1*, and *PICALM*. Additionally, the transcriptional start sites of EOAD genes *APP* and *PSEN1* also show significant H3K27ac enrichment in oligodendrocytes relative to other cell types. Previous multi-scale network analyses have found oligodendrocyte transcript and protein modules are enriched for genes associated with AD risk loci, particularly *BIN1* and *PICALM*^69,70^. Indeed, *BIN1* is highly expressed in oligodendrocytes, and is associated with white matter tracts in the human brain^71^. Combined, these data suggest epigenomic mechanisms in oligodendrocytes play a significant role in the functionality of certain AD risk loci and their associated risk genes^72^.

In parallel, we also find oligodendrocyte-enriched glia show by far the largest acetylation differences associated with Aβ load. In the hippocampus, the promoters of genes associated with early and late-onset AD risk displayed hypoacetylation. This includes EOAD risk genes *APP, PSEN1*, and *PSEN2*, and several genes associated with LOAD risk, including *BIN1, PICALM, ADAM10, ADAMTS4, FERMT2*, and *SORL1*^25–27,73^. Sorted hippocampal oligodendrocyte nuclei from an independent cohort of ROSMAP individuals were used to assess transcript levels of these genes, which revealed a corresponding downregulation of transcripts in individuals with high Aβ load. This suggests that EOAD and LOAD may share common pathogenic mechanisms in oligodendrocytes. In addition to risk genes, H3K27ac peaks associated with core oligodendrocyte processes such as myelination were significantly hypoacetylated in the hippocampus of AD subjects. Myelin-associated genes *STMN4* and *MYRF* were confirmed to have corresponding transcriptional downregulation in the same independent cohort of ROSMAP individuals.

The hippocampus is one of the earliest brain regions affected by AD pathology^74^. Here, we describe the first cell type-specific H3K27ac dataset from the hippocampus of postmortem AD patients. This provides a resource by which we can understand the epigenomic signatures of distinct cell types at a crucial anatomical locus of neurodegeneration. Specifically, it is evident that the H3K27ac changes in oligodendrocyte-enriched glia provide insight as to how the epigenomic state of the hippocampus is altered in AD. White matter lesions are positively correlated with hippocampal atrophy, and white matter hyperintensities are thought to be a core feature of AD^75,76^. Thus, the marked hypoacetylation observed near genes associated with Aβ processing and myelination in hippocampal oligodendrocytes suggest these biological processes are defective and may directly contribute to AD progression. Importantly, previous AD studies demonstrate similar pathways are deregulated at the transcriptomic and proteomic levels in oligodendrocyte-enriched modules, as does a recent single-cell gene expression study^64,69,70^. Combined with our current findings, this strongly suggests oligodendrocytes play an active role in AD progression and merit further attention. Although we identified these hypoacetylated peaks from female AD patients, the lack of sex-specificity observed in supporting publications and in our RT-qPCR validation lead us to conclude these DARs most strongly reflect epigenomic changes associated with Aβ load.

Interestingly, our dataset reveals dlPFC and hippocampus oligodendrocyte-enriched populations mount distinct epigenomic signatures in response to AD. Similar to our findings in the hippocampus, we observed an Aβ-correlated deregulation of myelin-associated promoters and enhancers in dlPFC oligodendrocytes. However, these dlPFC DARs become hyperacetylated in AD individuals, as do peaks annotated to *PSEN2, CLU, ADAMTS4, BIN1*, and *SORL1*. The DARs in the hippocampus are largely distinct from the DARs in the dlPFC, indicating brain region-specific epigenomic alterations. This disparity between brain regions may reflect oligodendrocyte heterogeneity in response to pathological insults, as well as region-specific differences in cell composition and pathologic severity. Alternatively, it may be associated with compensatory signaling in the prefrontal cortex that has been previously reported in neurodegenerative disorders^77^. However, in total, it is apparent oligodendrocyte H3K27ac represents a core feature of epigenomic dysregulation in both hippocampus and dlPFC.

Many lines of evidence have revealed roles for oligodendrocyte-driven myelination processes in both multiple sclerosis and major depression^78–80^, and studies are ongoing to advance the understanding of glial cells in neurological disorders^81^. The connection between AD and oligodendrocyte epigenomic dysregulation is not well-understood, and our data highlight this topic as a priority for future research. We propose further investigation into the role of myelination and demyelination is warranted in AD.

Our full set of DARs constitutes a larger list of novel genomic targets related to oligodendrocyte function, Aβ processing and oxidative phosphorylation that may be targeted using genome editing technologies such as CRISPR-Cas9^66^. For example, the promoter of the disintegrin and metalloprotease *ADAMTS18* displays the strongest hyperacetylation in the dlPFC of AD subjects, revealing it as a strong candidate for future therapeutics.

Lastly, we foresee that single nucleus level epigenomic assays for transposase accessible chromatin (snATAC-seq) can enable understanding of disease associated epigenomic deregulation and cell type heterogeneity in disease at a resolution that supersedes our study and previous studies. However, while active enhancers and promoters commonly lie in accessible chromatin regions, H3K27ac is a more robust indicator of active gene regulation, and therefore, our data resource can augment such future studies.

Taken together, our study shows the power of cell type-specific epigenomic profiling in identifying pathways and genomic loci that are differentially regulated in AD. We reveal new cell type-specific processes involved in AD which opens opportunities to ameliorate its harmful effects by targeting therapeutics to oligodendrocytes.

## METHODS

### Source of Brain Tissue and Pathologic Data

Biospecimens and data came from autopsied participants in one of two prospective clinical-pathologic cohort studies, the Religious Orders Study or Rush Memory and Aging Project (ROSMAP). Both studies were approved by an Institutional Review Board of Rush University Medical Center. All participants signed an informed consent, an Anatomical Gift Act, and a repository consent to all their data and biospecimens to be repurposed. Details of the studies have been previously reported^43^.

### Fluorescence-Activated Nuclei Sorting

Frozen dorsolateral prefrontal cortex and hippocampus samples were retrieved from -80°C storage and thawed on ice, then disrupted with a handheld homogenizer. Samples were fixed with 1% paraformaldehyde for 10 minutes at room temperature. Fixation was quenched with glycine for 5 minutes. Nuclei were isolated by dounce-homogenization followed by filtration through a 70uM cell strainer (cat no. 21008-952, VWR, Radnor PA). To immunotag cell type specific nuclei, anti-NeuN antibody conjugated to Alexa Fluor 488 (cat no. MAB377X, EMD Millipore, Burlington MA), and anti-PU.1 antibody conjugated to Alexa 647 (cat no. 2240S, Cell Signaling Technology, Danvers MA) were incubated with nuclei at 4°C for one hour and overnight, respectively. Samples were strained through a 40um filter (21008-949, VWR) and stained with DAPI (D9542, Sigma Aldrich, St. Louis MO) before flow cytometry. Fluorescence activated nuclei sorting was performed until at least 400,000 nuclei were collected for each cell type (NeuN+, Pu.1+, and NeuN-/Pu.1-) using the FACSAria (BD Biosciences, US).

### Chromatin Immunoprecipitation

Following sorting, chromatin was fragmented into 200-600 bp fragments using the Diagenode bioruptor. Fragmented samples were split equally into two tubes such that each tube contained an equivalent of chromatin from 200,000 nuclei. All ChIPs were carried out in duplicate. Samples were pre-cleared with BSA-blocked Protein A sepharose beads (cat no. GE17-0780-01, Sigma Aldrich) for four hours at 4°C. At this point, 1% input was collected and stored at -20°C. Chromatin was incubated with 2ug of Histone H3 (acetyl K27) antibody (cat no. ab4729, abcam, Cambridge UK) overnight at 4°C. Chromatin fragments bound to the antibody were pulled down with BSA-blocked Protein A sepharose beads for four hours at 4°C. To reduce non-specific binding, the bead-chromatin complex was washed four times with ice-cold RIPA buffer. Immunotagged chromatin was eluted from beads through shaking at 65°C for 15 minutes. Both 1% input and ChIP were de-crosslinked overnight in T50E10S1 buffer at 65°C. Reverse crosslinked chromatin was treated with RNase A and Proteinase K. DNA was purified using phenol-chloroform extraction. Following ethanol precipitation, samples were resuspended in 10 mM Tris-HCl buffer and stored at -20°C.

### ChIP-seq high-throughput sequencing

A portion of the sample was used to assess enrichment for cell-type specific H3K27ac peaks via qPCR. If the sample passed qPCR quality control, libraries were generated from the remaining sample. Library generation was performed using the KAPA Hyper Prep Kit (KK8504, Kapa Biosystems). After amplification and quantification, a portion of the library was used for a second qPCR to ensure enrichment of cell-type specific H3K27ac peaks. If the sample passed the second qPCR quality control, it was submitted to the MIT BioMicro Center for fragment analysis, followed by sequencing. The 40-bp single-end sequencing was performed using the Illumina HiSeq2000 platform according to standard operating procedures.

### Peak Calling, Quality Control and Read Counting

For peak calling, the AQUAS ChIP-Seq workflow (https://github.com/kundajelab/chipseq_pipeline) was used. To perform quality control, the two technical replicates for each sample were individually input to the AQUAS workflow to compute standard ENCODE quality metrics^44^ such as NSC, RSC, NRF, PBC1, PBC2, FRiP, replicate consistency etc. All samples that did not meet quality standards of (NSC>1.01, RSC>0.4, PBC1>0.4) were discarded at this point. The workflow uses Burrows-Wheeler alignment^82^, Samtools^83^ for processing alignments, MACS2^84^ for peak calling, and PICARD (http://broadinstitute.github.io/picard/) for removing PCR duplicates. Peak reproducibility is assessed by overlapping peaks across groups of sample replicates and pseudoreplicates using a method similar to irreproducible discovery rate (IDR)^85^ analysis. All analysis was performed on the hg19 reference genome.

Reproducible peaks were called on samples pooled by each separate group of samples defined by brain region, cell type population and presence or absence of Aβ load. The mergeBed^86^ utility was then used to merge the set of peaks across these pools. At this step, peaks that were less than 200 bp apart were merged together to account for local depletions in chromatin intensity profiles (“dips”)^87^. We propose this merged peak set as a reference set for peaks active in different brain cell types in the dlPFC of AD and non-AD subjects and use it in downstream analyses. The featureCounts^88^ package was used to count the read signal at these peaks for every ChIP-Seq experiment. This read count matrix was then used in downstream analysis for validation of sorting and for identifying differentially acetylated regions using DESeq2^45^. ROSMAP subject metadata were used as post-hoc covariates in the analysis.

### Cell type peak sets

We also generated reproducible peak sets for each cell type by assessing reproducibility across the two brain regions for subjects without Aβ load. This peak set was used to generate the browser visualization tracks at the loci containing the *INPP5D, BIN1* and *PICALM* genes (**Figure 2d, e and f**). Browser tracks for *INPP5D, BIN1* and *PICALM* were generated using the integrative genomics viewer (IGV)^89^ and pygenometracks^90^, and edited later.

Further, for each of the three cell type populations, we used the negative binomial model of DESeq2^45^ to identify subsets of differentially hyperacetylated peaks in the focal cell type population against the two non-focal populations from the full set of brain peaks (see **Peak Calling, QC and Read Counting**). Peaks were defined as differentially hyperacetylated if they displayed a positive log fold change and passed an adjusted p-value threshold of 0.05. A cell type background peak set was then created from these three sets of peaks using the mergeBed utility. This set of peaks was used in heritability enrichment analyses using permutation testing^35,50^ and stratified LD-score regression^47–49^.

### Sorting validation and identification of cell types by comparison to single nucleus RNA-seq clusters

The full set of merged peaks were annotated to their nearest genes using the annotatePeaks tool in HOMER^91^. Marker gene sets for 15 single nucleus RNA-Seq cell type clusters were downloaded from Habib et al^46^. For each single nucleus RNA-seq cluster, the set of H3K27ac peaks for which the closest gene was present in the marker gene set was obtained. Then, DESeq2^45^ was used to compute log2FC at these peaks between ChIP-Seq samples corresponding to a focal foreground cell type population against ChIP-seq samples corresponding to the other two background cell type populations. A one-sided t-test was used to test whether the distribution of log2FC was significantly greater than 0.5 (∼1.4 fold change). A significant result from this test indicated the enrichment of a cell type in the focal ChIP-Seq population. The test was conducted for every pair of focal ChIP-Seq population and single nucleus RNA-Seq cluster. p-values were adjusted for multiple hypothesis testing using Bonferroni’s correction.

In addition, a similar approach was used to verify these results at the individual sample level as well as groups of samples defined by Aβ load, sex and brain region. Variance stabilized counts were used and mean log2FC for each focal population was computed against the other two non-focal populations. For each of the 15 cell type clusters, the mean log2FC was then computed for peaks annotated to that cluster and the resulting values were plotted in a heatmap.

To test whether distant peaks confound these results, the above analyses were also conducted on peaks that are near promoters of the marker genes by only considering the peaks that are less than 5 kilobases away from transcription start sites of the 15 gene sets.

### Enrichment test for colocalization of AD-associated variants with cell type-specific peaks

GWAS summary statistics from two studies, Kunkle et al^26^ and Jansen et al^27^ were downloaded. and stratified LD-score regression (S-LDSC)^47–49^ was used to compute AD SNP heritability in enrichment in differential peaks for each cell type against the merged background set. The standard workflow described by the authors was used and LD scores were computed based on custom annotations derived from hyperacetylated peaks called on each cell type and compared against custom annotations derived from the merged background set constructed from the three cell type hyperacetylated peak sets. The regression coefficients for each population were extracted and plotted. A significant result from this test indicates an enrichment of genetic risk for LOAD in regions that are actively regulating gene expression in the cell type, suggesting a role for that cell type in influencing predisposition towards LOAD.

To test whether choice of computational method may confound these results, we used another approach that utilizes a permutation test^35,50^. LD-pruning was applied (LD > 0.5) on both GWAS datasets based on the 1000 genomes reference^92^. SNPs overlapping protein coding sequence^93^ were filtered out along with SNPs in tight linkage disequilibrium (LD > 0.5). SNPs with p-values less than 1e-3 were selected and overlap annotations were created for each set of differential cell type-specific peaks (see **Cell type peak sets**). A permutation test was used to compute heritability enrichment of AD-associated SNPs in a focal foreground set of differential peaks for a cell type against the merged background set. SNPs were permuted 1,000,000 times preserving distance to gene, minor allele frequency and the number of variants that are in LD.

### DARs associated with Aβ load

Differentially acetylated regions were identified using the negative binomial model of DESeq2 on the previously generated count matrix (see **Peak Calling, QC and Read Counting)** selecting peaks associated with a binary Aβ load indicator. An adjusted p-value cutoff of 0.05 was used for selecting differentially acetylated peaks. For each differential acetylation model setting **(Supplementary Table 3)**, a reduced count matrix was generated that includes only the subset of samples corresponding to the variables described. Variance stabilized (vst) read counts^45^ across all peaks were used for heatmap visualization and principal components analysis (PCA). Box plots of read counts against covariate variables such as Aβ, age, years of education and sample quality metrics were produced using peak-wise Z-scores of vst normalized counts. DAR sets were annotated to their nearest genes using the annotatePeaks tool in HOMER^91^ and the distribution of distance to TSS was plotted for the two biggest DAR sets as well as the remaining DARs.

Genome browser visualizations were created for the two biggest DAR sets at known EOAD and LOAD risk loci, as well as highly differentially acetylated loci using pygenometracks^90^. Custom UNIX commands and the UCSC bigWigMerge^94^ tool were used to create average H3K27ac signal tracks in oligodendrocyte enriched glia samples of subjects with and without Aβ load. Tracks for DESeq2 log2FC and UCSC known gene annotations^95^ were included. Generated visualizations were edited later.

The Genomic Region Enrichment and Annotation Tool (GREAT)^59^ web tool was used for computing enrichments for ontological annotations associated with genes proximal to DAR sets. GREAT analysis was performed separately on the two biggest DAR sets as well as the remaining DARs not in those sets. In addition, we used GREAT to annotate neuron, microglia and oligodendrocyte enriched glial hyperacetylated peaks for enriched functions. The merged brain peak set (see **Peak Calling, QC and Read Counting**) was used as the background for each GREAT analysis. A heatmap of the fold enrichment returned by GREAT was plotted for any GO Biological Process that passed a q-value cutoff of 0.05 and was associated with a minimum of 5 genes in any of the 6 GREAT analyses. In addition, fold enrichment for the KEGG Alzheimer’s Disease Pathway was plotted in the heatmap.

S-LDSC was used to test for AD SNP heritability enrichment from both AD GWAS studies in the two biggest DAR sets. The full brain peak set was used as background.

### Correlation based clustering of DARs

Variance stabilized (vst) counts for all Aβ associated DARs were obtained. All samples passing quality control were included. Clustering was performed separately for TSS proximal (<5kb) DARs and TSS distal (>5kb) DARs. A distance matrix based on Pearson correlation was computed between every pair of peaks. More specifically, the distance between peak *x* and peak *y* was calculated as *1-abs*(*cor*(*x,y*)), where *abs* represents absolute value and *cor* represents Pearson correlation. The absolute value was used because it gives equal weightage to negatively correlated peaks and positively correlated peaks. Then, average linkage hierarchical clustering using the hclust R function was performed using this distance matrix to construct a dendrogram. To identify stable clusters from the resulting dendrogram, the a dynamic tree cutting approach^96^ was used. The resulting cluster identities were plotted alongside a heatmap of variance stabilized counts.

### RNA extraction, reverse transcription and quantitative PCR in postmortem hippocampus (qPCR)

An independent set of hippocampal samples from the ROSMAP cohort were used for rt-qPCR validation. Samples were prepped for FANS as described previously. To isolate oligodendrocyte, microglia, astrocyte, and neuronal nuclei, samples were stained overnight at 4°C with anti-Olig2 antibody conjugated to Alexa Fluor 488 (cat no. MABN50A4, EMD Millipore, Burlington MA), anti-PU.1 antibody conjugated to Alexa Fluor 647 (cat no. 2240S, Cell Signaling Technology, Danvers MA), anti-GFAP conjugated to Alexa Fluor 555 (cat no. 3656, Cell Signaling Technology, Danvers MA), and stained for one hour with anti-NeuN conjugated to biotin (cat no. MAB377B, EMD Millipore, Burlington MA), and for one hour with Brilliant Violet 711 Streptavadin (cat no. 405241, BioLegend, San Diego, CA). Fluorescence activated nuclei was performed until at least 100,000 Olig2-positive nuclei, NeuN-positive nuclei, GFAP-positive nuclei, and PU.1-positive nuclei were collected for each sample.

Following sorting, nuclei were treated for 15 minutes with Proteinase K at 50°C and then for 13 minutes at 80°C. RNA was extracted using Direct-zol RNA MicroPrep kit (Zymo Research) according to manufacturer’s instructions. Reverse transcription of RNA was carried out using Invitrogen SuperScript IV First Strand Synthesis System (Oligo dT) according to manufacturer’s protocol. qPCR was performed using a Bio-Rad CFX-96 quantitative thermocycler and SsoFast EvaGreen Supermix (Bio-Rad). Relative changes in gene expression were determined using the 2^-ΔΔCt^ method. The geometric mean of cycle numbers from RPL13, CYC1, and GADPH were used for housekeeping Ct values. Primer sequences used for qPCR can be found in Supplementary Table 6.

### Comparison with snRNA-seq from postmortem dlPFC

Hyperacetylated DARs in oligodendrocyte enriched glia of the dlPFC were assessed for nearby transcriptional differences identified in the snRNA-seq study from Mathys, Valderrain et al^64^. The nearest genes of the hyperacetylated DARs were obtained using annotatePeaks in HOMER. The oligodendrocyte cluster specific log2FC of these genes was obtained from the snRNA-seq study. Then, a one-sample one-sided t-test was used to test whether there is an average increase in transcription at these genes (null hypothesis log2FC = 0, alternative hypothesis log2FC > 0). Transcriptional fold-change of specific AD risk genes and genes near highly hyperacetylated peaks was also plotted.

### DARs associated with Age

Age associated changes in H3K27ac levels were identified using DESeq2. To control for potential confounds, sex, the binary Aβ load status, and brain region were added as covariates in the linear model along with age. An adjusted p-value cutoff of 0.05 was then used to screen for peaks differentially acetylated with every unit increase in age. The peaks were grouped as having increased or decreased acetylation with age, and ontological annotation enrichments were computed using GREAT (see **DARs associated with Aβ load**) using a full brain background peak set. For heatmap visualization, variance stabilizing transformation (vst) was applied on the full matrix and differential peaks were extracted.

## DATA AVAILABILITY

The ChIP-seq data will be made available on The Rush Alzheimer’s Disease Center (RADC) Research Resource Sharing Hub at https://www.radc.rush.edu/docs/omics.htm or at Synapse (link to be provided) under a doi. The ROSMAP metadata will be accessible at (link to be provided). The data will be available under controlled use conditions set by human privacy regulations. To access the data, a data use agreement will be needed. This registration is in place solely to ensure anonymity of the ROSMAP study participants. A data use agreement will be agreed with either Rush University Medical Center (RUMC) or with SAGE, who maintains Synapse, and will be downloadable from their websites. Bed, narrowPeak and bigwig files that do not contain private information will be made available at the appropriate resource in accordance with privacy considerations.

## CODE AVAILABILITY

Code for processing and analyzing the data will be made available at: https://github.com/pfenninglab/ad_h3k27ac_3ct

## AUTHOR INFORMATION

These authors contributed equally: Easwaran Ramamurthy and Gwyneth Welch

These authors jointly supervised the work: Li-Huei Tsai and Andreas Pfenning

## CONTRIBUTIONS

J.C., G.W., E.R., D.A.B., A.P. and L.-H.T., all contributed to study design. The study was coordinated and directed by A.P. and L.-H.T. J.C. performed FANS and ChIP-seq for dorsolateral prefrontal cortex samples. G.W. performed FANS, ChIP-seq and RT-qPCR for hippocampus samples. E.R. performed ChIP-seq processing and led the computational analyses. Y.Y. did the aging analysis supervised by A.P. and E.R. L.G. performed initial genetics integration analysis supervised by A.P.. E.R., G.W., J.C., Y.Y., D.A.B, L-H.T., and A.P wrote and edited the manuscript.

## ACKNOWLEDGEMENTS

We thank the study participants and staff of the Rush Alzheimer’s Disease Center. This work was supported in part by the Cure Alzheimer’s Fund (CAF) and The Okawa Foundation for Information and Telecommunications (A.P.); NIH grants RO1NS102730, RO1AG054012, and The Glenn Foundation for Aging Research (L.-H.T.). E.R was supported by a Presidential fellowship from Carnegie Mellon Brainhub. G.W. was supported by the Barbara Weedon Champions of the Brain Fellowship from MIT Brain and Cognitive Sciences department.

## ETHICS DECLARATION

A.P served as a paid consultant for Cognition Therapeutics, Inc during preparation of the manuscript.

## SUPPLEMENTARY FIGURES

**Supplementary Figure 1:**
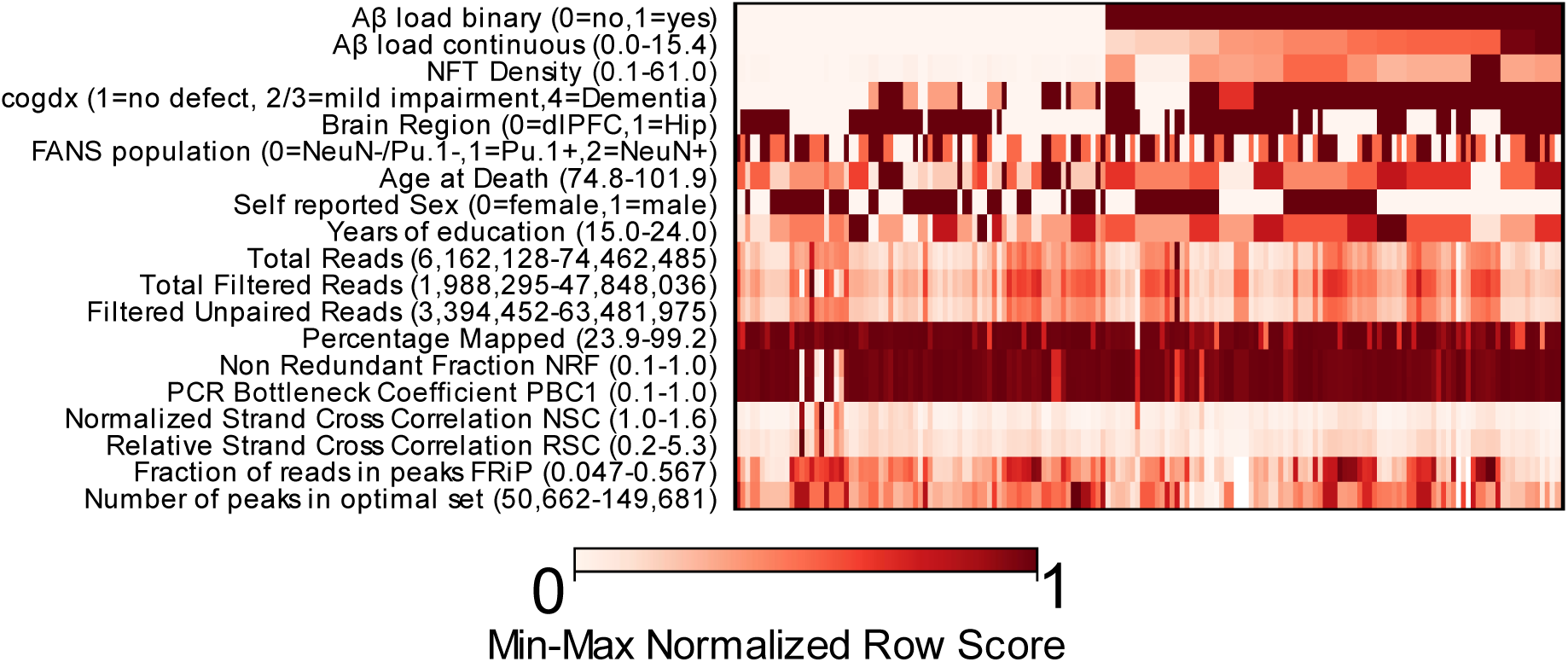
Heatmap of collected subject information and standard ENCODE quality metrics for all ChIP-Seq samples

**Supplementary Figure 2:**
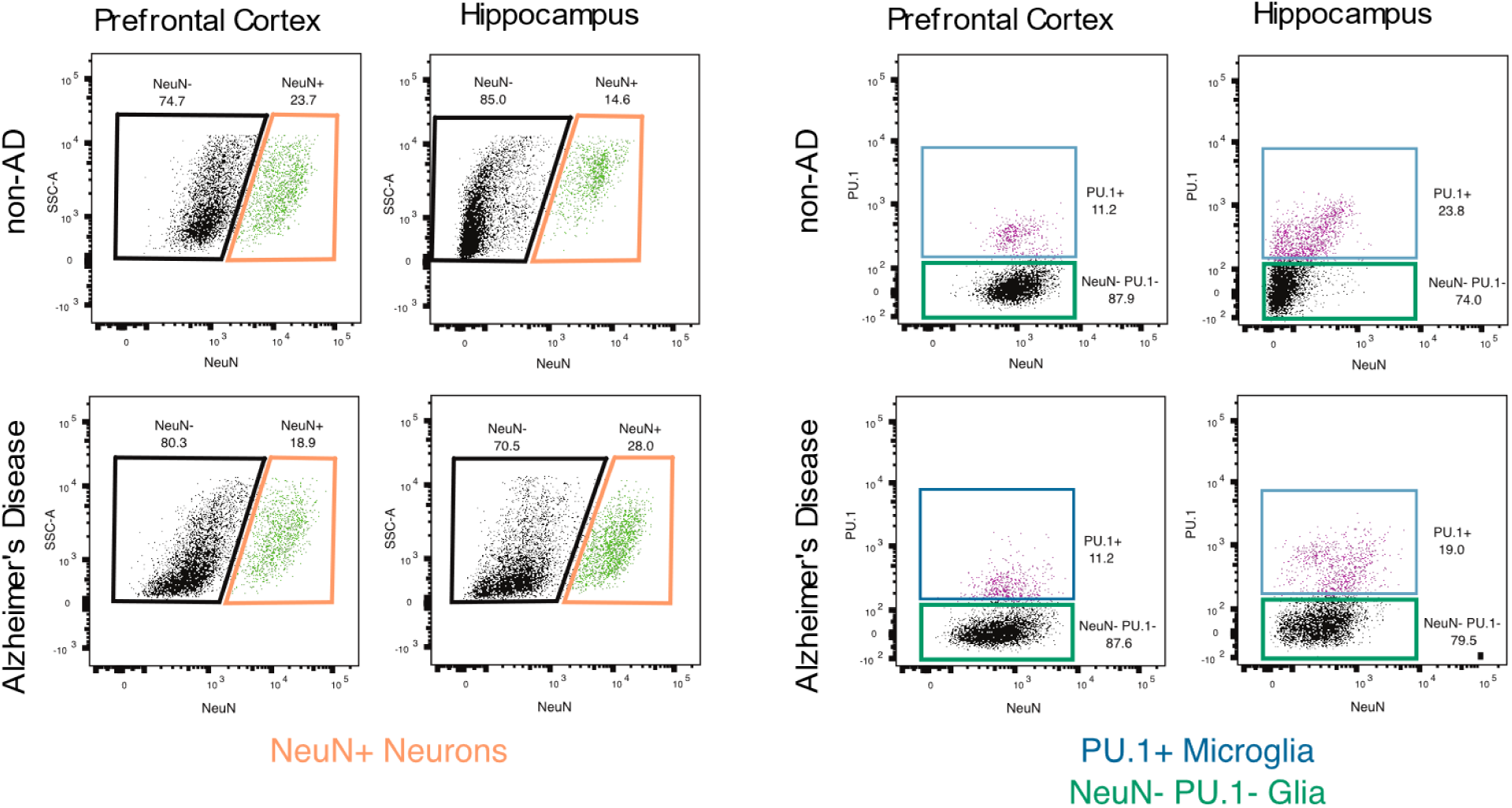
Sorting of neuronal and microglial nuclei from prefrontal cortex and hippocampus of subjects with and without AD

**Supplementary Figure 3:**
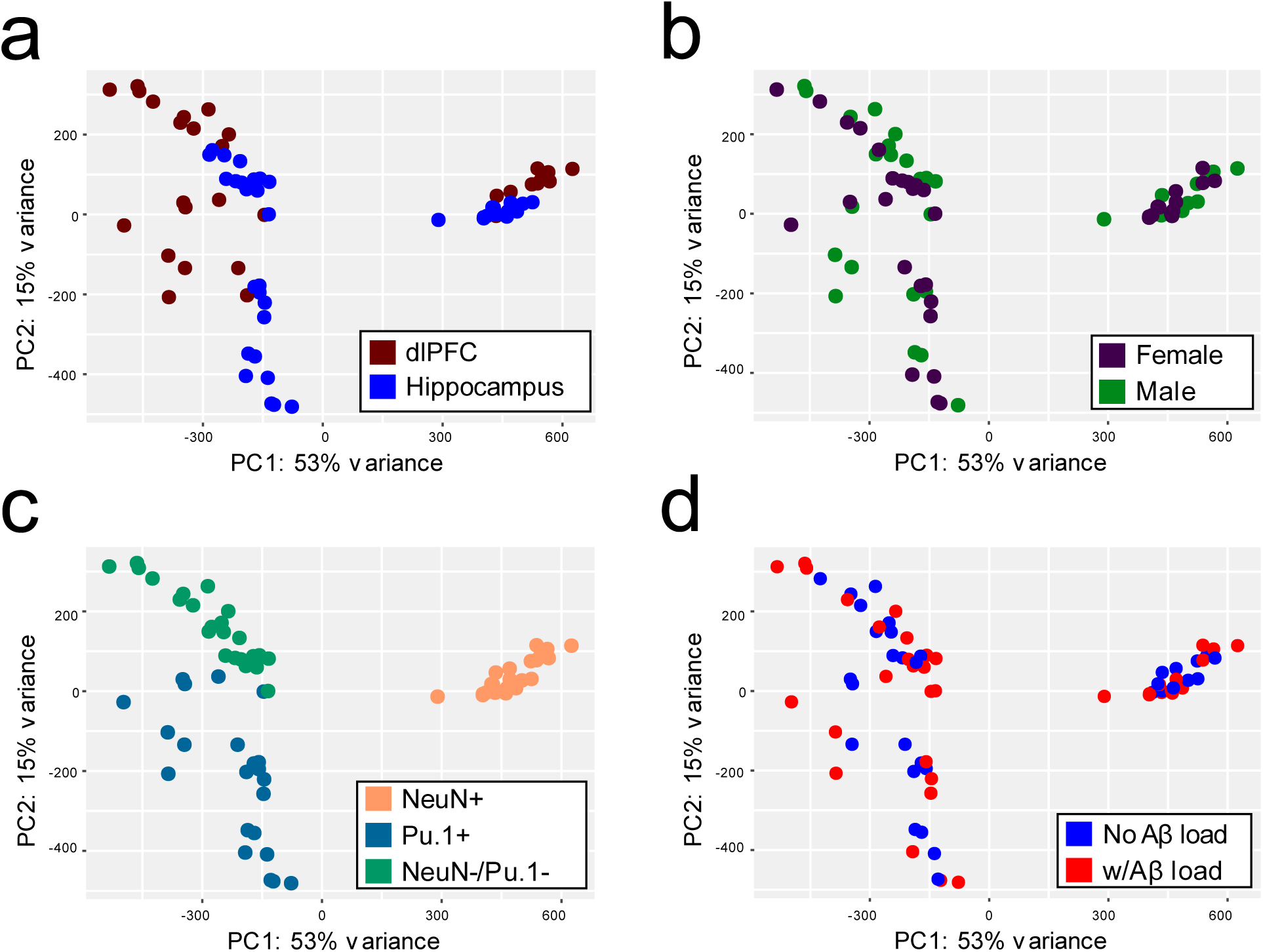
Plots showing all sequencing samples projected into principal components space using PCA colored by a. brain region b. sex c. cell type population d. Aβ load. PC1 separates NeuN+ samples from Pu.1+ and NeuN-/Pu.1-samples and PC2 separates Pu.1+ samples from NeuN-/Pu.1-samples

**Supplementary Figure 4:**
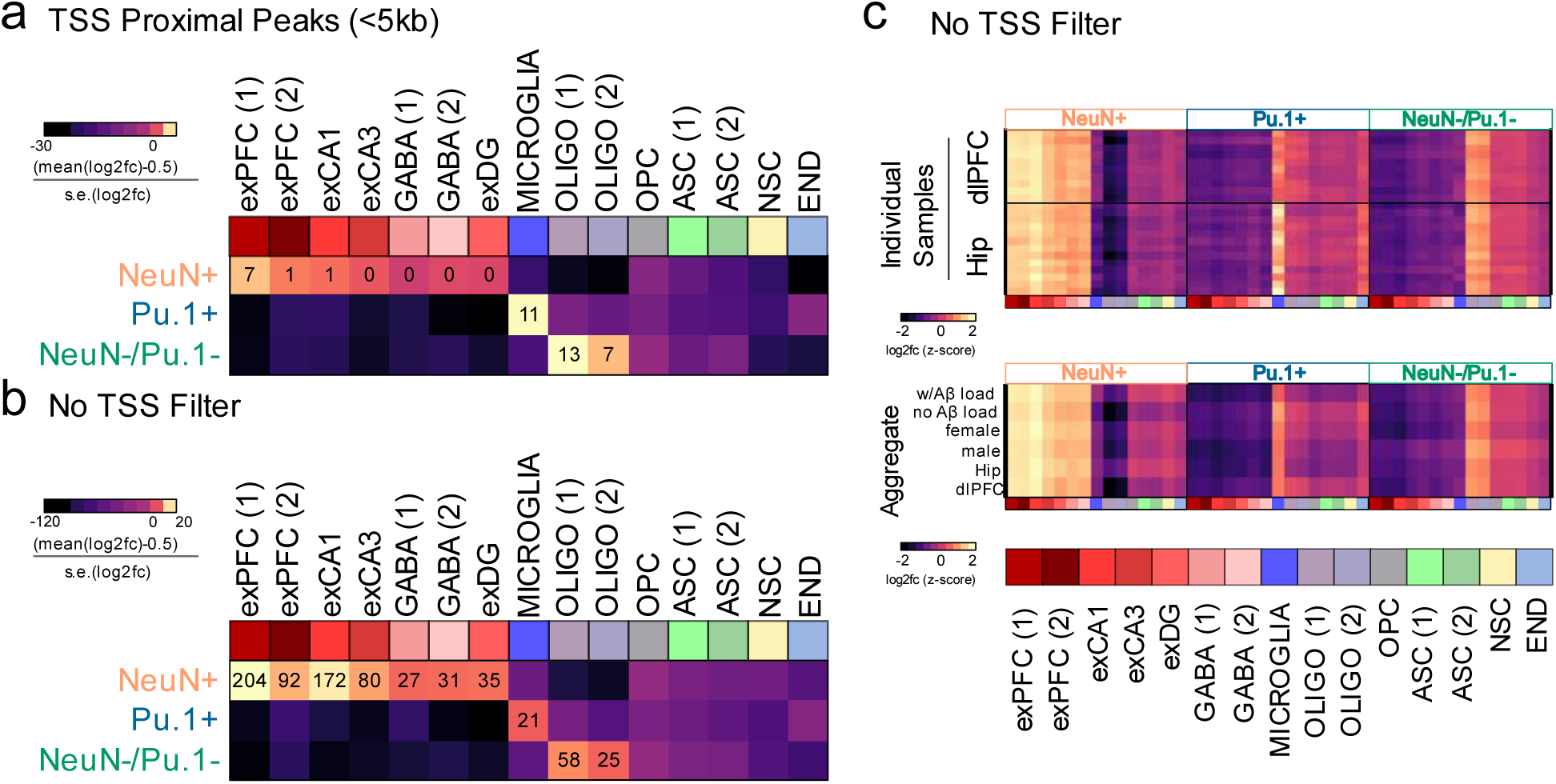
Analysis of cell type compositions of FANS populations using H3K27ac signal at peaks near 15 cell type-specific gene lists annotated in Habib et al^46^. **a**. t-statistic showing enrichment of a given cell type cluster in a FANS population over the other two non-focal FANS populations. Mean log2FC of signal was computed at H3K27ac peaks near the promoters (<5kb distance to transcription start site) of the 15 cell type-specific marker gene sets and a t-test was used to compute whether the mean log2FC is greater than 0.5 (∼1.4-fold change). Value inside box represent p-value (-log10-transformed) for the t-test **b**. same as panel a but using no distance to tss filter for defining the cell type-specific peaks **c**. mean log2fc (focal population/non-focal populations) for peaks near 15 cluster markers at the individual sample level and for sample aggregates using no distance to tss filter for defining the cell type-specific peaks. Abbreviated labels for the single nucleus analysis clusters are presented. exPFC, glutamatergic neurons from the PFC; GABA, GABAergic interneurons; exCA1/3, pyramidal neurons from the hippocampus CA region; exDG, granule neurons from the hippocampus dentate gyrus region; ASC, astrocytes; MICROGLIA, microglia; OLIGO, oligodendrocytes; OPC, oligodendrocyte precursor cells; NSC, neuronal stem cells; END, endothelial cells.

**Supplementary Figure 5:**
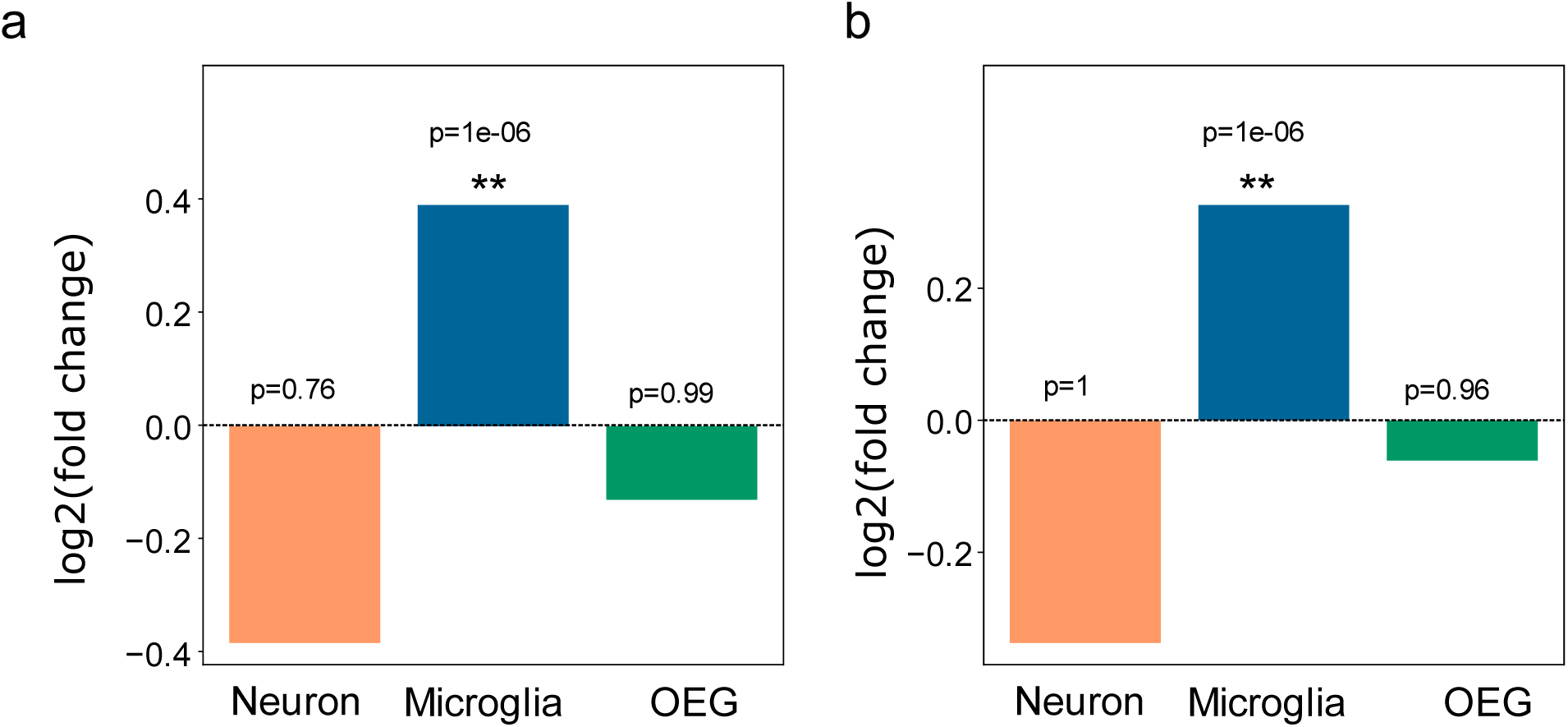
Independent permutation test method confirms that AD associated variants from the two large GWAS studies^26,27^ colocalize with peaks enriched in the microglial population relative to peaks enriched in the oligodendrocyte and neuronal population suggesting that microglial gene regulation may influence predisposition towards LOAD. **a**. plot of fold change (log2 transformed) in the number of AD associated SNPs (Kunkle et al. GWAS p-value < 1e-3) overlapping focal foreground cell type specific peaks versus the number of AD associated SNPs overlapping background set of peaks in all three cell types; permutation test controls for LD and minor allele frequency, p-value for colocalization test is indicated above each bar **b**. same as **a**. but using the Jansen et al GWAS.

**Supplementary Figure 6:**
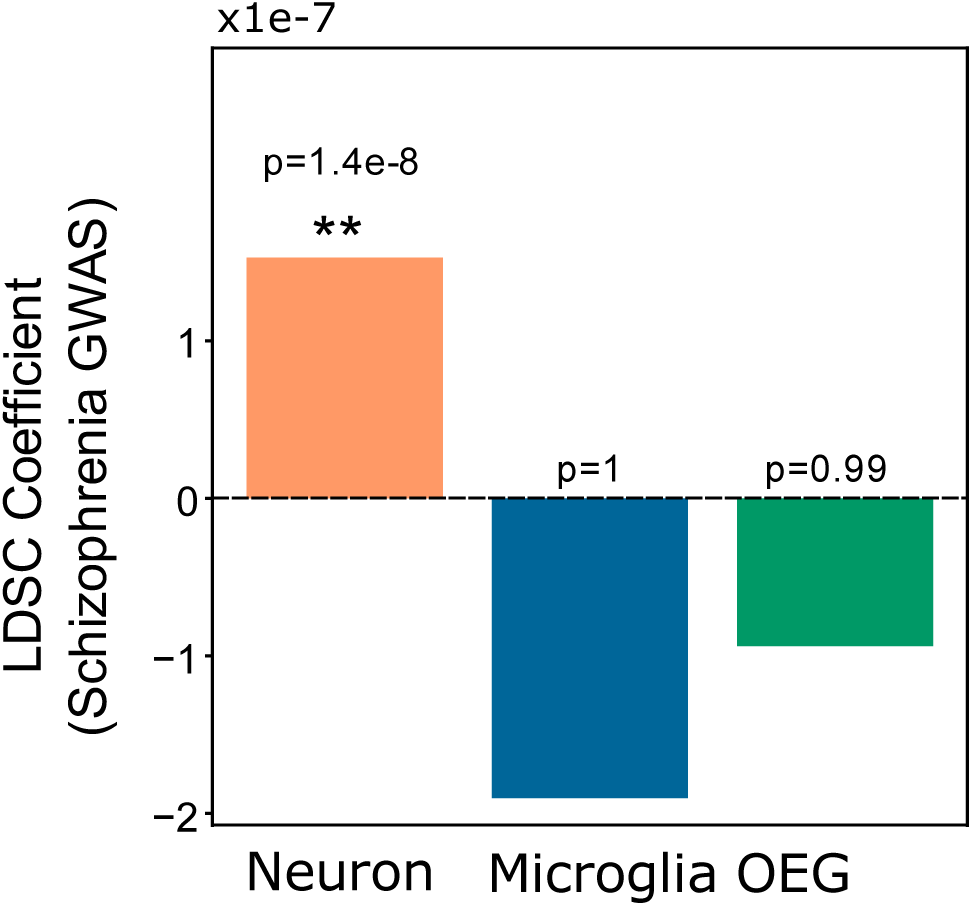
SNPs associated with Schizophrenia from a large GWAS^53^ colocalize with peaks enriched in the neuronal population relative to peaks enriched in the OEG and microglial population suggesting that neuronal gene regulation may influence predisposition towards Schizophrenia. Plot shows the estimated stratified LD score regression coefficient for the three peak sets. p-values are indicated above each bar.

**Supplementary Figure 7:**
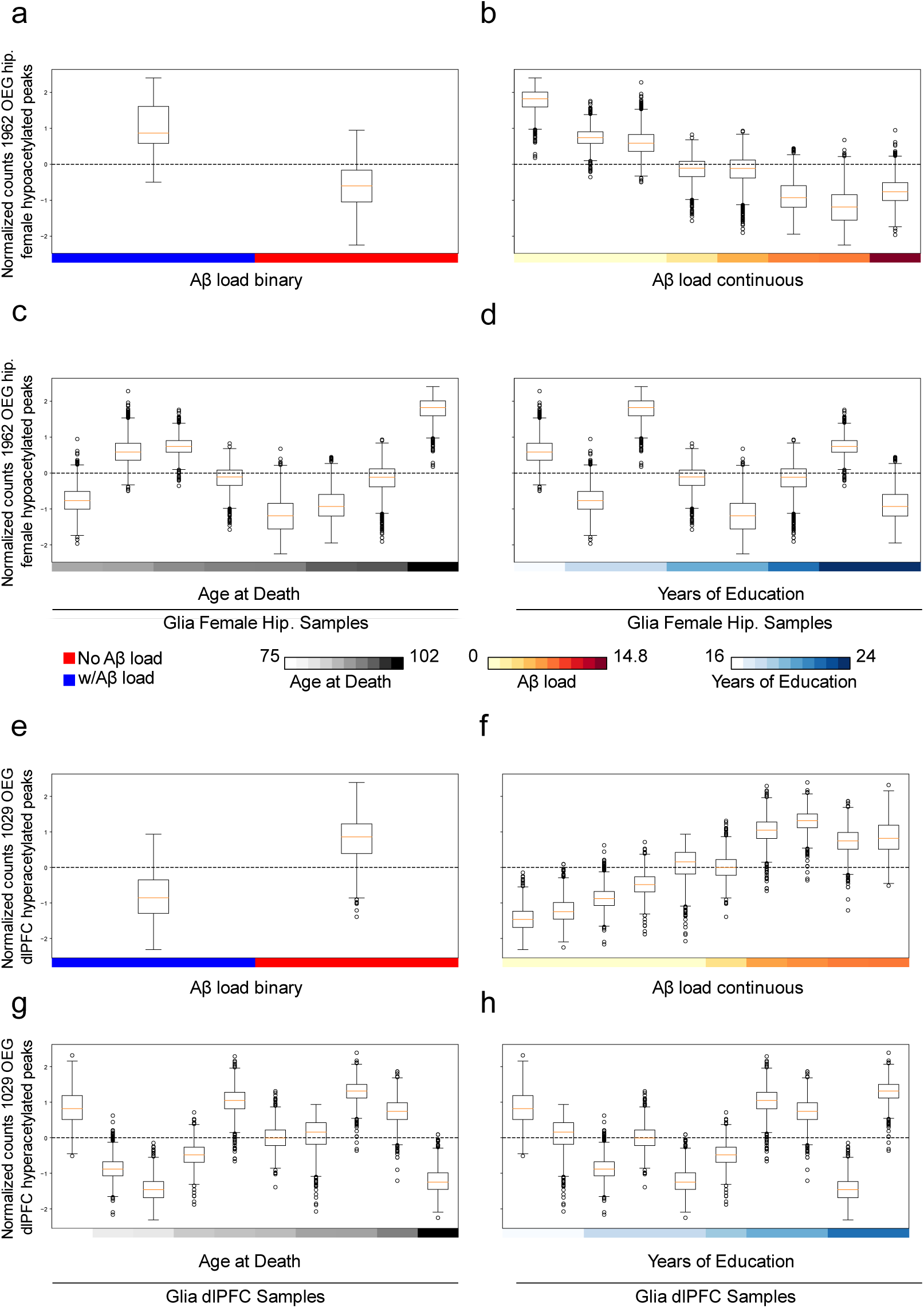
Two largest DAR sets of 1962 hypoacetylated peaks in female hippocampus OEG and 1029 hyperacetylated peaks in dlPFC OEG do not show correlations with other covariates such as age at death and years of education. **a and b**. Read counts at 1962 hypoacetylated DARs show consistent decrease with an increase in overall Aβ **c and d**. No relation can be observed between mean read count and variables such as age at death and years of education **d**,**e**,**f and g**. same as a,b,c and d but for 1029 hyperacetylated DARs in dlPFC OEG

**Supplementary Figure 8:**
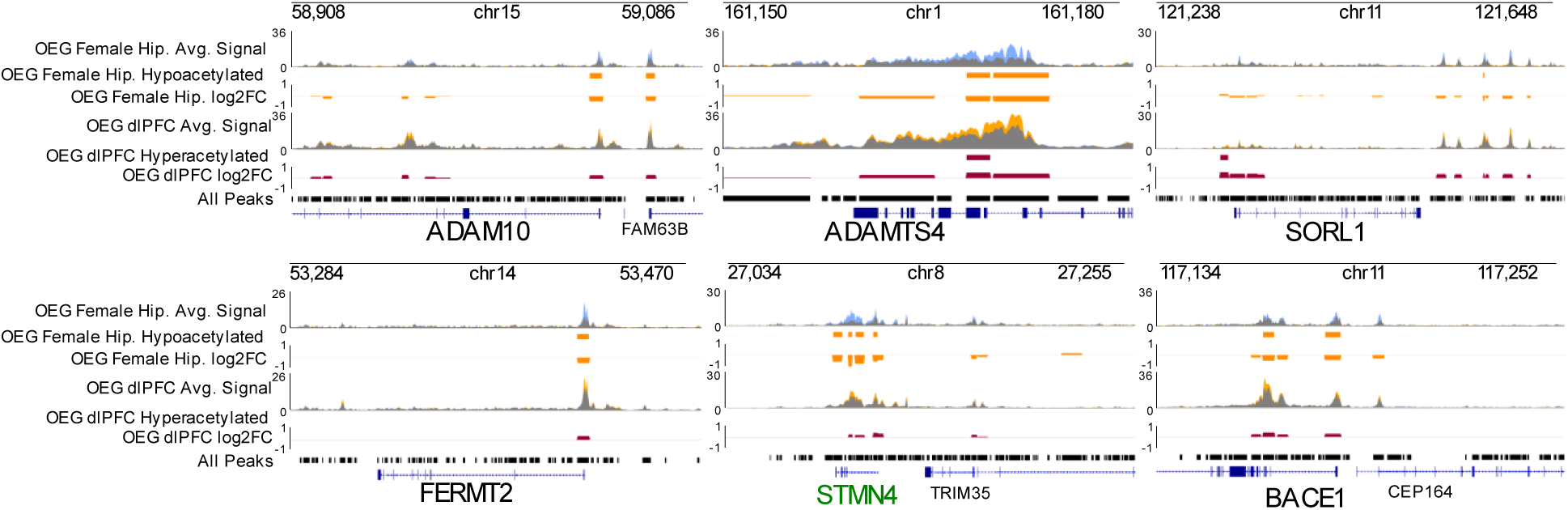
Genome browser tracks displaying average H3K27ac signal in OEG samples of subjects with (yellow) and without Aβ load (blue) corresponding to the two biggest DAR sets at known LOAD risk loci, as well as strongly differentially acetylated regions near *STMN4*

**Supplementary Figure 9:**
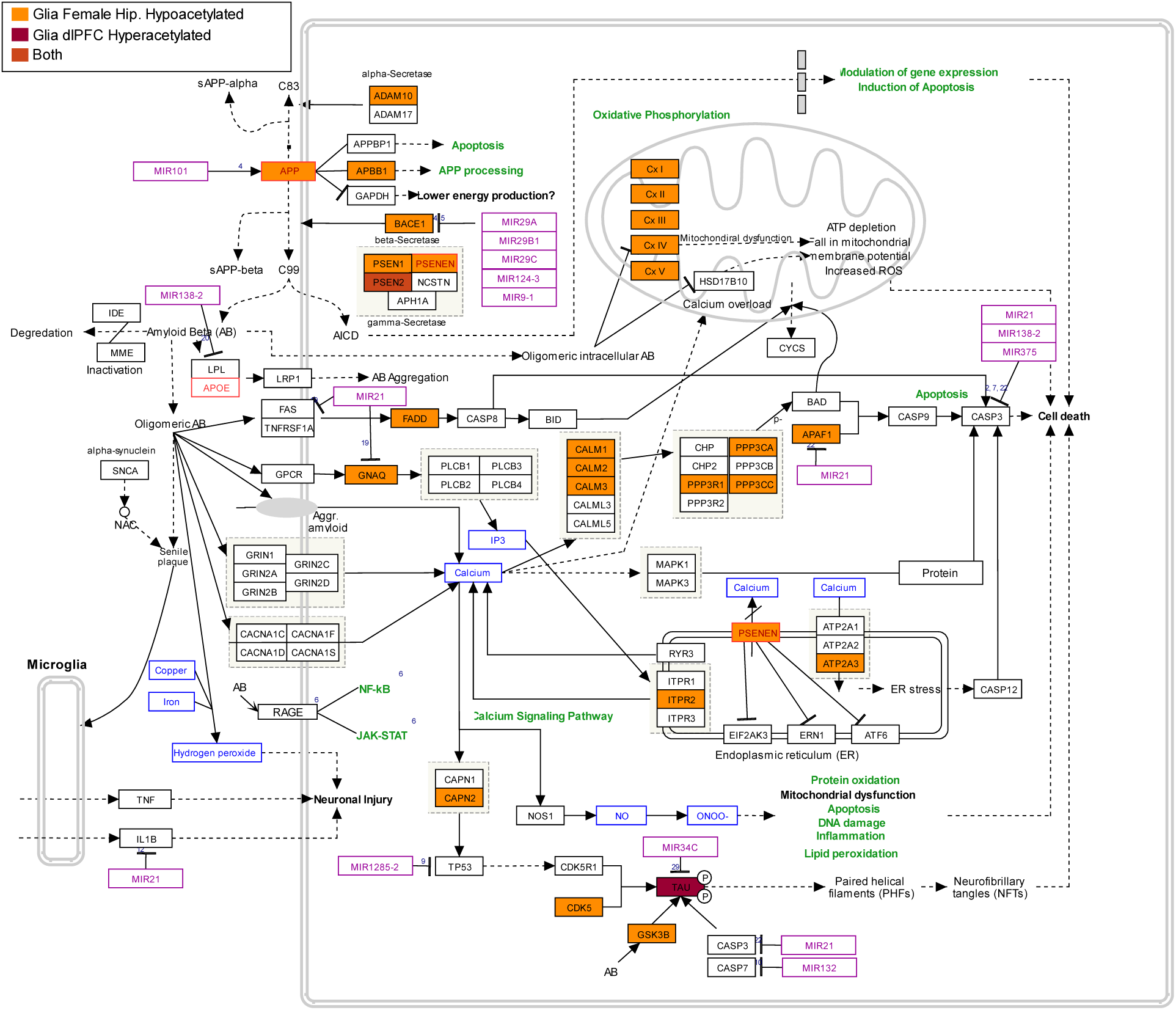
Peaks associated with genes in multiple modules in the KEGG Alzheimer’s Disease Pathway are part of the two largest OEG specific DAR sets and include peaks near all three of the secretase complexes, *MAPT* and all 5 modules involved in oxidative phosphorylation. Boxes colored in orange represent genes which contain an annotation to the 1962 hypoacetylated peaks discovered in female hippocampus OEG, boxes colored in maroon represent genes which contain an annotation to the 1029 hyperacetylated peaks in dlPFC OEG, an average of orange and maroon is used to color boxes which represent genes present in both sets

**Supplementary Figure 10:**
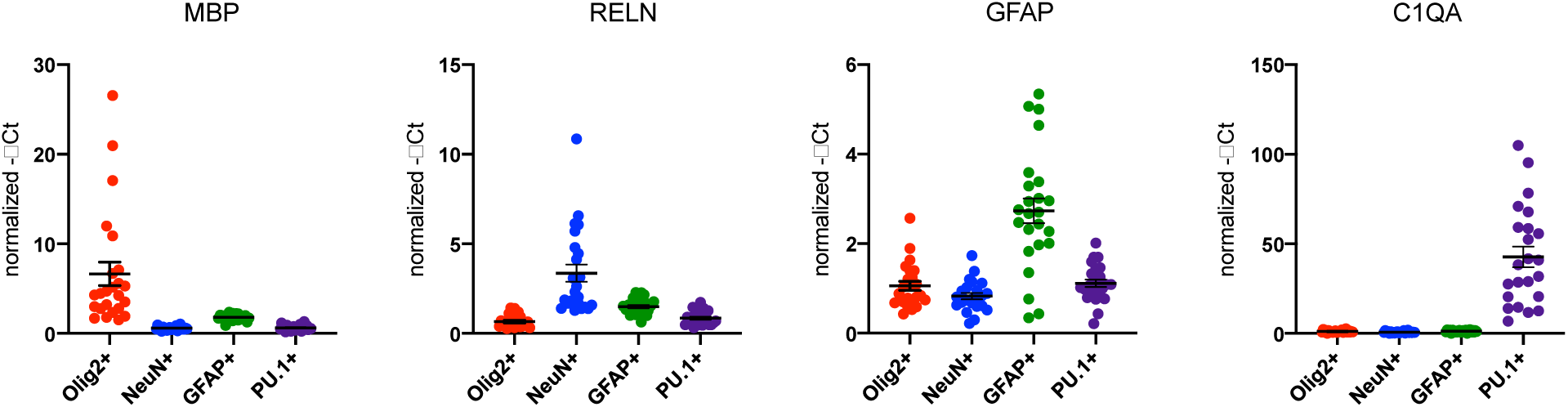
mRNA from cell type specific nuclei are enriched for cell type specific markers. Olig2+ nuclei are enriched for oligodendrocyte-specific mRNA MBP, NeuN+ nuclei are enriched for neuron-specific mRNA Reln, GFAP+ nuclei are enriched for astrocyte-specific mRNA GFAP, and Pu.1+ nuclei are enriched for microglia-specific mRNA C1qa. S.E.M. displayed.

**Supplementary Figure 11:**
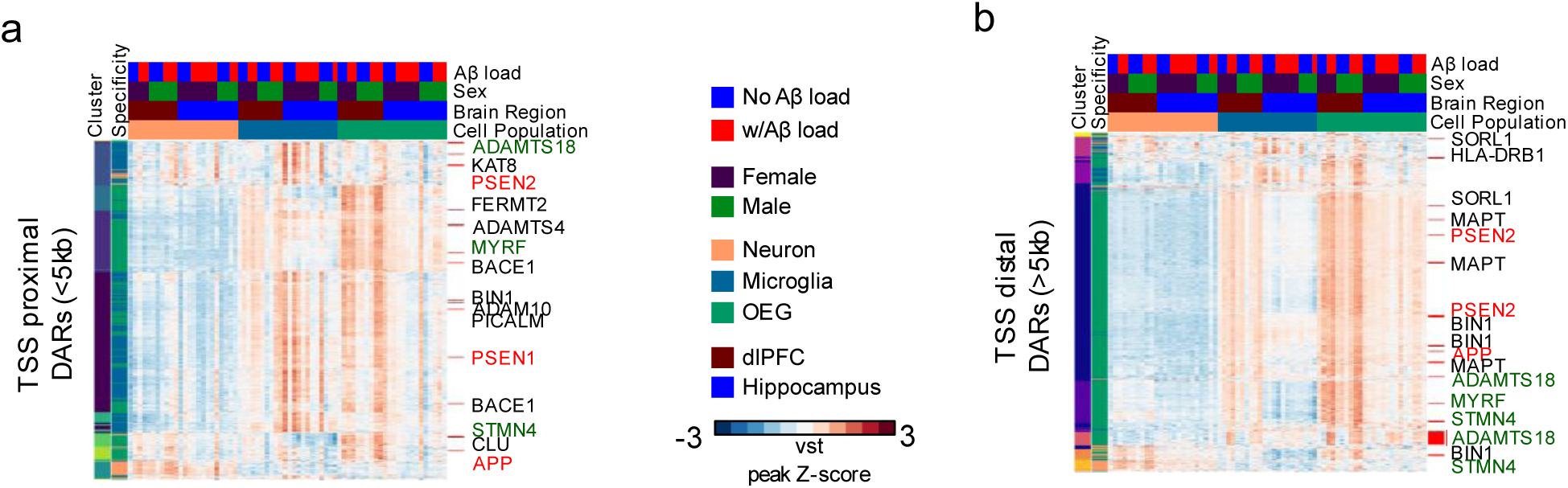
**a**. correlation-based clustering of all TSS proximal DARs identified across different sex, brain region and cell type populations. Heatmap displays normalized acetylation levels at each of the DARs across all samples for every cell type and brain region. Cell type-specificity and cluster membership are indicated for each peak on the left of the heatmap **b**. same as **a** but for TSS distal DARs.

**Supplementary Figure 12:**
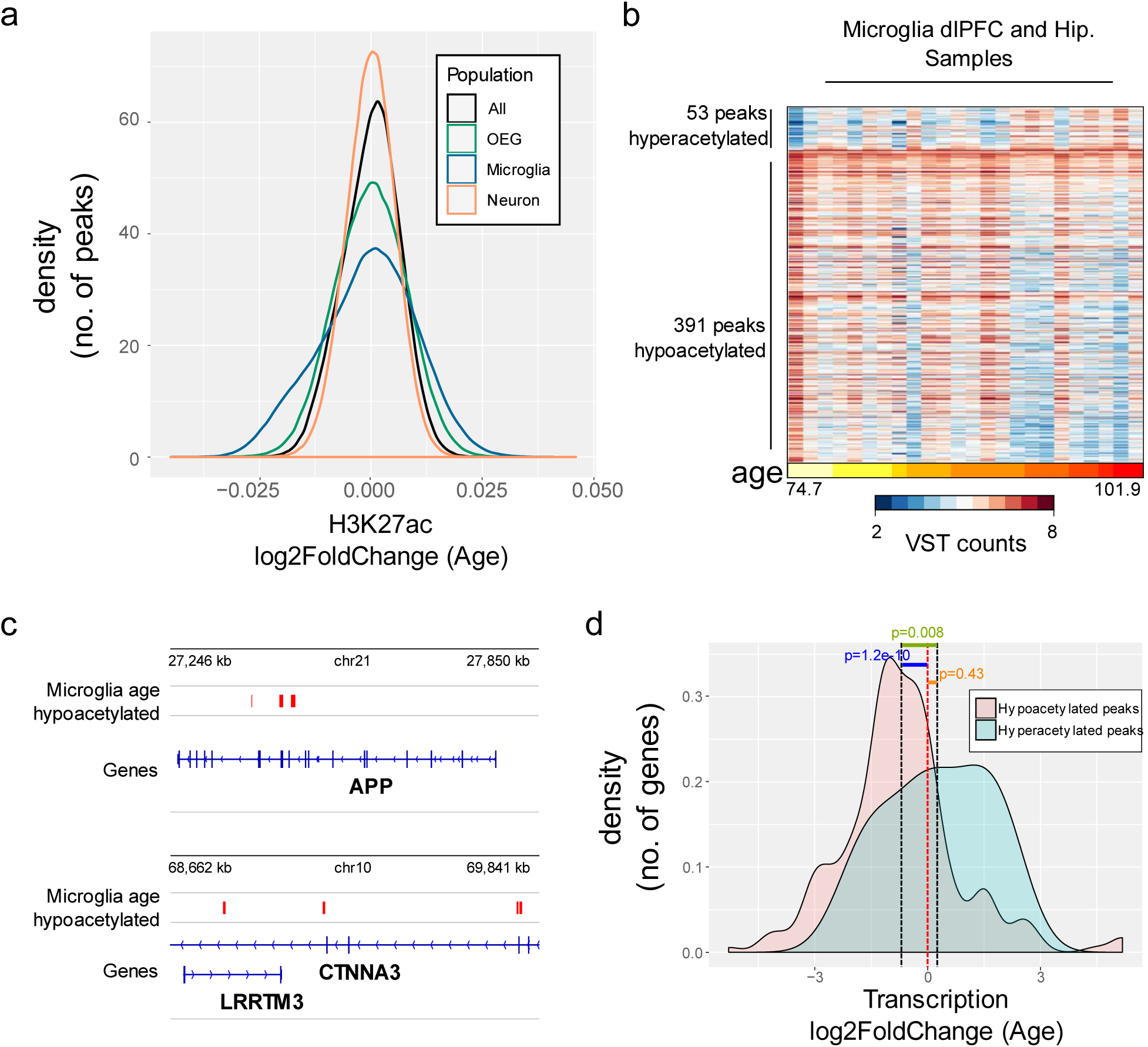
Summary of peaks displaying differential H3K27ac with age in microglia. **a**. distribution of age associated log2 fold changes for the entire peak set shows that age associated changes in H3K27ac levels are enriched in the microglial population - as indicated by the wider distribution of log2 fold changes relative to total, neuronal, and OEG populations, **b**. heatmap of variance stabilized read counts representing acetylation levels for the 444 peaks that display age associated differential H3K27ac levels in the microglial population, **c**. Genome browser tracks displaying age hyperacetylated peaks near *APP* and *LRRTM3* which are both involved in Aβ processing **d**. distribution of fold change (log2 transformed) in transcription identified in Olah et al^97^ for genes annotated to 391 age hypoacetylated peaks and 53 age hyperacetylated peaks. Two dashed black vertical lines represent mean transcription log2FC for genes near age hypoacetylated and age hyperacetylated peaks, respectively. Red vertical dashed line represents log2FC=0, representing no difference in transcription. T-test p-values are indicated for three comparisons, from left to right (i) hypoacetylated mean vs. 0, (ii) hyperacetylated vs hypoacetylated means, and (iii) hyperacetylated mean vs. 0

